# CRNKL1 is a highly selective regulator of intron-retaining HIV-1 and cellular mRNAs

**DOI:** 10.1101/2020.02.04.934927

**Authors:** Han Xiao, Emanuel Wyler, Miha Milek, Bastian Grewe, Philipp Kirchner, Arif Ekici, Ana Beatriz Oliveira Villela Silva, Doris Jungnickl, Markus Landthaler, Armin Ensser, Klaus Überla

**Author notes:** Corresponding author: Prof. Dr. Klaus Überla, Institute of Clinical and Molecular Virology, University Hospital Erlangen, Friedrich-Alexander Universität Erlangen-Nürnberg, Schlossgarten 4, 91054 Erlangen, Germany. these two authors contributed equally (Emanuel Wyler, Miha Milek).

## Abstract

The HIV-1 Rev protein is a nuclear export factor for unspliced and incompletely-spliced HIV-1 RNAs. Without Rev, these intron-retaining RNAs are trapped in the nucleus. A genome-wide screen identified nine proteins of the spliceosome which all enhanced expression from the HIV-1 unspliced RNA after CRISPR/Cas knock-down. Depletion of DHX38, WDR70 and four proteins of the Prp19-associated complex (ISY1, BUD31, XAB2, CRNKL1) resulted in a more than 20-fold enhancement of unspliced HIV-1 RNA levels in the cytoplasm. Targeting of CRNKL1, DHX38, and BUD31 affected nuclear export efficiencies of the HIV-1 unspliced RNA to a much larger extent than splicing. Transcriptomic analyses further revealed that CRNKL1 also suppresses cytoplasmic levels of cellular mRNAs with selectively retained introns. Thus, CRNKL1 dependent nuclear retention seems to be a novel mechanism for the regulation of cytoplasmic levels of intron-retaining cellular mRNAs that is harnessed by HIV-1 to direct its complex splicing pattern.

## Introduction

Intron retention is increasingly recognized as a regulatory mechanism of gene expression and diversification of the cellular proteome (1). Transcriptomic analyses indicate that a large percentage of eukaryotic genes encode transcripts retaining introns (2, 3). Intron-retaining transcripts may be trapped in the nucleus leading to their degradation. Alternatively, specialized nuclear export pathways may enable certain intron-retaining transcripts to reach the cytoplasm, where they can be translated or undergo nonsense-mediated decay. Retroviruses use intron retention to encode multiple proteins from one primary transcript. RNA-secondary structures or cellular or viral adaptor proteins link intron-retaining retroviral transcripts in the nucleus to the TAP/NXF1 or the CRM-1 nuclear export factors (4), leading to the export of intron-retaining RNAs. While a number of studies explored how nuclear retention can be overcome, the factors mediating nuclear retention of intron-containing mRNAs are mostly unknown, but are likely linked to partial assembly of splicing complexes on intron-containing RNA (5).

The first evidence for a regulated export and expression of intron-retaining RNAs in mammalian cells was obtained in studies investigating the replication of human immunodeficiency virus type 1 (HIV-1). Alternative splicing and intron retention leads to three types of transcripts: fully-spliced (FS) mRNAs, incompletely-spliced (IS) mRNAs, and the unspliced (US) genomic RNA (6). The process of viral gene expression is temporally regulated, resulting in early expression of regulatory proteins and late expression of structural proteins (7, 8). In the early phase, the US and IS transcripts are retained in the nucleus, and are subjected to the nuclear splicing or degradation machinery (9). Only FS transcripts are transported to the cytoplasm via the default mRNA export pathway, TAP/NXF1, leading to expression of Tat, Rev and Nef proteins (10, 11). The regulatory proteins Tat and Rev are nuclear cytoplasmic shuttle proteins and regulate viral gene expression. While Tat transactivates transcription by enhancing RNA elongation, Rev fine-tunes posttranscriptional processes and acts as a switch to initiate the late phase of the viral replication cycle (reviewed in Grewe and Uberla, 2010; Mousseau and Valente, 2017). It binds to the Rev-responsive element (RRE) which is present in the HIV-1 US and IS transcripts, but not in the FS transcripts, and multimerizes on the RRE. The protein exporting factor CRM1 and associated factors are recruited by the Rev-RRE complex, leading to the nuclear-cytoplasmic export of these two classes of RNA (14).

Nuclear retention of intron-retaining HIV-1 RNAs has been explained by their suboptimal splice sites, leading to the trapping of the intron-retaining RNAs in the spliceosome and subsequent degradation (15). In addition, multiple inhibitory sequences (INS) were identified in the retained introns (16–20). The presumed inactivation of these inhibitory sequences by codon-optimizing the AT-rich HIV-1 sequence abolished nuclear retention (21) independent of the presence or absence of HIV-1 splice donor and splice acceptor sites (22, 23). Why AT-rich intron-containing HIV-1 sequences, but not the codon-optimized variants are retained remains elusive. It has been reported that knock-down of the paraspeckle lncRNA NEAT1 could promote HIV Gag production through increased nucleocytoplasmic export of INS-containing RNAs, implicating the nuclear paraspeckles in the nuclear retention of HIV US and IS RNAs (24). However, this has been questioned by a study demonstrating that HIV-1 US RNA was not actually colocalized with paraspeckles (25), despite that several paraspeckle proteins (PSF, Martin3 and RBM14) have been reported to associate with HIV-1 INS elements and/or promote the Rev-dependent RNA export (26–28).

In addition to these viral determinants and lncRNA contributing to nuclear mRNA retention, depletion of hnRNPA2/B1 has been shown to increase cytoplasmic levels of HIV-1 genomic RNA, but this increase did not enhance Gag expression levels and the precise mechanism remains unknown (29). With regard to nuclear retention of mRNA in general, a cellular complex named pre-mRNA retention and splicing complex (RES) has been identified in yeast (30). The RES is composed of Bud13, Snu17p, and Pml1P and inactivation of either of these genes in yeast reduces splicing of an intron with a weak 5’splice donor. In addition, Pml1P inactivation enhanced expression of a pre-mRNA reporter construct without substantial reduction of the spliced mRNA suggesting a role of Pml1P and the RES in nuclear retention. More recently, BUD13 was identified to bind to the poorly-spliced mRNA of mammalian IRF7, a master regulator of the interferon response (31). Knock-down of BUD13 specifically enhanced retention of intron 4 of IRF7 in stimulated cells thus reducing IRF7 protein levels and dampening the interferon response. However, intron-containing IRF7 mRNA could not be detected in the cytoplasm (31), indicating additional nuclear trapping mechanisms or rapid cytoplasmic degradation of intron-containing IRF7 mRNA. HIV-1 introns also differ from typical RES-dependent introns (31–33) by their low GC content, challenging the hypothesis that HIV-1 splicing may be regulated by the RES.

An alternative nuclear trapping mechanism could be a cellular quality control mechanism for mRNAs located at the nuclear pore complex (NPC). Experimental or stress-induced interference with this complex results in leakage of intron-retaining mRNAs into the cytoplasm (34–37). This finding suggests that the NPC-associated quality control complex also has a role in nuclear retention of intron-containing mRNAs, probably as a back-up mechanism not influencing the initial splicing events. Here we identified cellular proteins involved in nuclear trapping of intron-retaining HIV-1 mRNAs by performing a genome-wide screen for cellular factors, whose depletion resulted in enhanced expression from HIV-1 unspliced transcripts in the absence of its export factor Rev. Furthermore, we examined the effect of identified cellular factors on HIV US and FS RNAs localization and expression. This revealed that CRNKL1 is a nuclear retention factor of the HIV-1 unspliced RNA and a highly selective regulator of cytoplasmic levels of intron-retaining HIV-1 and cellular mRNAs.

## Results

### Reporter cell lines for HIV-1 Rev-independent expression from the unspliced RNA

The overall strategy to identify cellular proteins involved in the nuclear trapping of intron-retaining HIV-1 mRNAs was to inactivate cellular genes by a genome-wide lentiviral CRISPR-Cas knock-out library and to select knock-out cells that show enhanced Gag-expression from a proviral Rev-deficient HIV-1 reporter construct. To easily quantify HIV-Gag expression in living cells, its ORF was fused to the blue-fluorescent protein (BFP) gene. The *rev* gene was inactivated by two point mutations and the DsRed reporter gene was expressed in place of *nef* from a fully-spliced HIV-1 transcript in order to mark HIV infected cells in the absence of Rev (Fig. 1A). Since we aimed to establish stable cell clones containing such proviral reporter constructs, expression of *pol, vif, vpr, and vpu* was also blocked by point mutations to avoid difficulties due to potential cytotoxic or cytostatic effects of these viral proteins (38–40). The *env* gene was inactivated by a 4-bp deletion resulting in a frame shift. Although this reporter construct, designated HIV-dual-GT only encodes the <u>G</u>ag-BPF fusion protein, DsRed, and <u>T</u>at (for sufficient transcriptional activation), all known *cis*-acting sequences potentially interacting with cellular components should be maintained.

**Figure 1.**
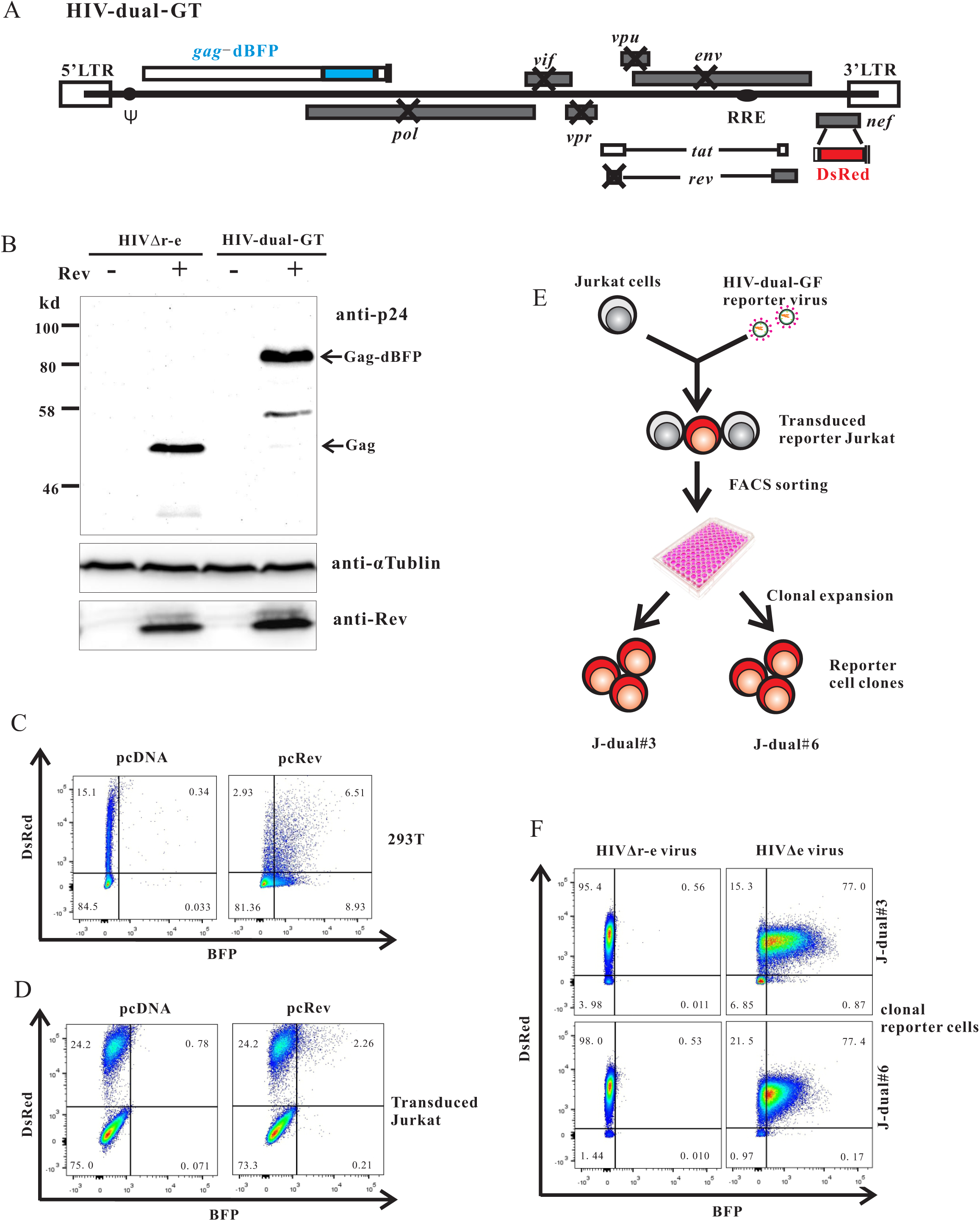
Establishment of HIV-dual-GT reporter cell clones. (A) Map of the HIV-dual-GT proviral reporter construct. (B) Western blot analysis of lysates of HEK293T cells transfected with HIV-dual-GT or HIVΔr-e in the presence or absence of an Rev expression plasmid (pcRev) with antibodies to the p24 capsid protein, αTublin or Rev. (C) 293T cells transfected with HIV-dual-GT and pcRev or pcDNA3.1 were analyzed by flow cytometry for Gag-BFP and DsRed expression. (D) Jurkat cells transduced with the HIV-dual-GT reporter vector (MOI 0.3) and transiently transfected with pcRev or pcDNA control plasmid were analyzed by flow cytometry. (E) Scheme for the generation of reporter cell clones. (F) HIV-dual-GT-transduced Jurkat reporter cell clones J-dual#3 and J-dual#6 were superinfected with VSV-G-pseudotyped HIV-1 viruses only differing in the expression of Rev and analyzed by flow cytometry. All assays were performed at least twice and results of one representative experiment are shown.

To characterize the expression pattern of HIV-dual-GT, HEK293T cells were transfected with plasmids containing HIV-dual-GT or a *rev*-deficient *env*-deletion mutant of HIV in the presence or absence of a Rev-expression plasmid. Western blot analyses revealed Rev-dependent expression of the Gag-BFP fusion protein (Fig. 1B). Flow cytometric analysis of HEK293T cells transfected with HIV-dual-GT revealed a major DsRed positive, but BFP negative cell population, that shifted to a double positive population by co-transfection of a Rev-expression plasmid (Fig. 1C). DsRed expression seemed to decrease in the presence of Rev consistent with reduced splicing and consequently reduced generation of fully-spliced DsRed mRNA due to rapid export of unspliced transcripts (7).

To generate reporter cell lines harboring a properly integrated proviral DNA of HIV-dual-GT, it was encapsidated and pseudotyped by cotransfection with HIV-gag-pol and VSV-G expression plasmids. The Rev deficiency was also complemented by cotransfection of a Rev-expression plasmid. Titers of the HIV-dual-GT vector were in the range of 5×10^5^ TU (transforming units)/ml. Flow cytometric analysis of Jurkat cells transduced with this VSV-G pseudotyped HIV-dual-GT vector particles revealed a distinct population of cells that was DsRed positive, but BFP negative (Fig. 1D). Transient transfection of this transduced bulk culture with a Rev-expression plasmid shifted approximately 10% of the DsRed positive population to a DsRed and BFP double-positive population. The magnitude of this effect was lower than in the co-transfection experiments in 293T cells (Fig. 1C) consistent with the lower transfection efficiency for Jurkat cells.

To obtain more homogenous reporter cells, Jurkat cell clones containing HIV-dual-GT were established by transducing Jurkat cells at an MOI of 0.1, followed by flow cytometric sorting of single DsRed positive, BFP negative cells into wells of a 96-well plate (Fig. 1E). Two of the expanded DsRed positive cell clones, showing a rather homogenous population of DsRed positive and BPF-negative cells and designated J-dual#3 and J-dual#6, were transduced at an MOI of 2 with VSV-G peudotyped *env* deletion mutants of HIV containing or lacking a functional *rev* gene. In the presence of a functional *rev* gene the BFP fluorescence was strongly induced in both reporter cell clones (Fig. 1F). Therefore, both clones were selected for the screening for cellular genes that suppress BFP expression from the HIV-dual-GT reporter virus.

### Genome-wide screening for cellular factors suppressing expression from intron-containing HIV-1 genomic RNA

Poor expression levels of HIV-1 structural genes in the absence of Rev that can be overcome by modifying the codon-usage, suggest that there is an active cellular suppressive mechanism acting on the level of the viral transcript. To identify cellular proteins necessary for this postulated suppressive mechanism, we used a GeCKOv2 genome-wide CRISPR-Cas knock-out library kindly provided by the laboratory of Feng Zhang (41, 42). The plasmid library contains 122,411 different single guide RNA genes (sgRNA) targeting 19,050 human genes (6 sgRNAs per gene) and 1,864 sgRNA genes targeting miRNA genes (4 sgRNAs per miRNA gene), and includes 1,000 non-targeting (NT) sgRNA genes. Cas9, sgRNA and puromycin-resistance genes are expressed from a single lentiviral vector construct, lentiCRISPRv2. After amplification of the plasmid library and confirmation of its diversity by next generation sequencing (data not shown), a stock of a VSV-G pseudotyped lentiviral vector library was prepared by transient transfection of 293T cells and titrated on the Jurkat reporter cell clones, J-dual#3 and J-dual#6, and parental Jurkat cells. All three cell lines were then transduced with the lentiviral CRISPR-library at an MOI of around 2. Four days after transduction, the 0.01 to 0.02% DsRed positive cells showing the highest BFP expression levels were sorted for next generation sequencing of the sgRNA genes delivered to these cells by the lentiviral vector library. To be able to determine enrichment of sgRNA genes enhancing Gag-BFP expression levels, the representation of each sgRNA gene from non-selected Jurkat cells transduced with the lentiviral CRISPR-library was also determined. Selective enrichment of sgRNA sequences in sorted J-dual#3 and J-dual#6 cells compared to non-sorted Jurkat cells was then determined by MAGeCK, a computational tool developed for the analysis of CRISPR screens (43, 44). The calculated robust ranking aggregation (RRA) score reflects enrichment of sgRNA sequences targeting the same gene in the selected cells. The distribution of the RRA score for all targeted genes is plotted in Fig. 2C with the top 12 candidate genes highlighted. For eleven of these, at least two different sgRNA sequences targeting the same gene were enriched in the selected cells arguing against confounding off-target effects of single sgRNA (Fig. 2D). The probability of false discovery of the top 12 candidate genes ranged from 0.0012 to 0.15.

**Figure 2.**
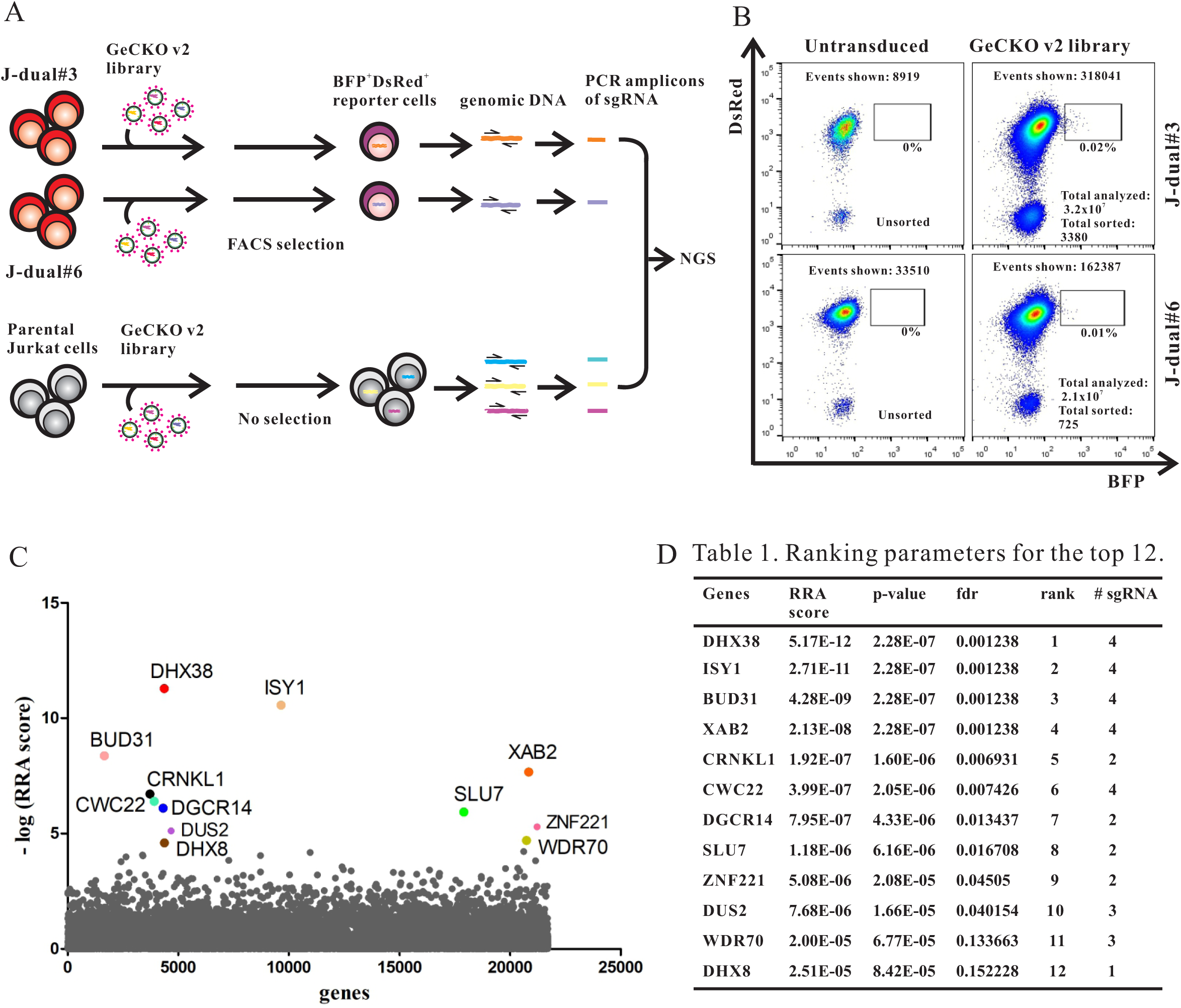
Genome-wide CRISPR/Cas screen. (A) Outline of screening strategy. (B) Flow cytometric analysis of J-dual#3 and J-dual#6 reporter cells without transduction and four days after transduction by the GeCKO v2 lentiviral vector library (MOI around 2) immediately prior to the sorting process. Total number of events shown in the graph, total number of cells analyzed in the screen and total numbers of cells sorted are indicated. (C) Enrichment analysis for guide RNAs in transduced and selected J-dual#3 and J-dual#6 reporter cells compared to transduced but unselected Jurkat cells by the MAGeck program. The robust ranking aggregation (RRA) score reflects the enrichment of guide RNA targeting the same gene in cells selected from both reporter cell clones. The distribution of the negative log10 of the RRA score for all targeted genes in alphabetical order on the X-axis is shown with the top 12 candidate genes highlighted. (D) Additional MAGeck ranking parameters of the top 12 candidate genes. fdr, false discovery rate; # sgRNA, number of enriched guide RNAs targeting the same gene.

### Analyzing interaction of validated hits shows a network of pre-mRNA splicing factors

Based on the enrichment of the individual sgRNAs targeting each of the top 12 candidate genes in the Gag-BFP positive cells selected by flow cytometry (Fig. S1), two sgRNA sequences were chosen for each of the candidate genes (Table S1A) for further validation. Gene-specific sgRNA sequences and a non-targeting sequence, named NT1, were cloned individually into the lentiCRISPRv2 vector plasmid. J-dual#3 and J-dual#6 reporter cells were then transduced with lentiviral vector particles transferring these sgRNA sequences. Upregulation of BFP expression was observed by flow cytometry for 10 of the 12 candidate genes. For nine of them, transfer of both sgRNA sequences upregulated BFP expression in both reporter cell clones (Fig. S2A). For DUS2 and ZNF221 the screening results could not be confirmed. Since long half-lifes of mRNAs and proteins may prolong the time from inactivation of the targeted gene to the decline of protein levels, we continued to monitor the reporter cells transduced with sgRNA genes targeting DUS2 and ZNF221 for up to 2 weeks post infection. However, upregulation of Gag-BFP remained undetectable (data not shown). In addition, we confirmed sufficient transduction rates by the lentiviral DUS2 and ZNF221 targeting vectors, excluding the possibility that reduced lentiviral vector titers are responsible for their failure to upregulate BFP expression (Fig. S2B). Despite this, the entire screening approach was highly specific showing a false discovery rate of only 0.166.

A first hint on potential mechanisms by which inactivation of the identified cellular genes could enhance Gag-BFP expression in the reporter cell lines was obtained by searching for functional protein association networks between the identified genes using the STRING online database (https://string-db.org/). Strikingly, 9 of the 10 confirmed hits are involved in pre-mRNA splicing (Fig. 3). Among them is, ESS2/DGCR14 for which no interactions with the other identified factors have been reported, but which is associated with U6 snRNA as well as U1 and U4 snRNAs (45). WDR70 is not linked to splicing, however, it contains a WD40 repeat domain which is shared by a number of splicing factors (46). The pre-mRNA splicing process includes two excision-steps catalyzed by a large and dynamic RNP machine. The confirmed hits DHX38, DHX8, SLU7 and DGCR14 are typical second-step factors. In contrast, XAB2, ISY1, CRNKL1, CWC22 and BUD31 should be recruited at the first step, and intriguingly all belong to the nuclear PPR19-associated complex (47–50). This complex is involved in diverse nuclear functions such as DNA double strand break (DSB) repair, transcriptional elongation, and splicing (51). The core factor of the complex, PRPF19 (human Prp19), however, was not identified in the screen. Consistently, targeting PRPF19 by two sgRNAs in J-dual#3 did not affect Gag-BFP expression (Fig. S3A). Since most of the confirmed hits are components of the spliceosome, we analyzed whether PRPF8, a ATPase/helicase at the core of the spliceosome (52), could also regulate the HIV Gag expression. Moreover, BUD13 from the RES complex and Tpr from the NPC, which were reported to have a nuclear retention function, were tested as well by using two different CRISPR/Cas sgRNAs targeting each of the genes. None of these targeting sgRNAs tested induced BFP expression (Fig. S3B-D).

**Figure 3.**
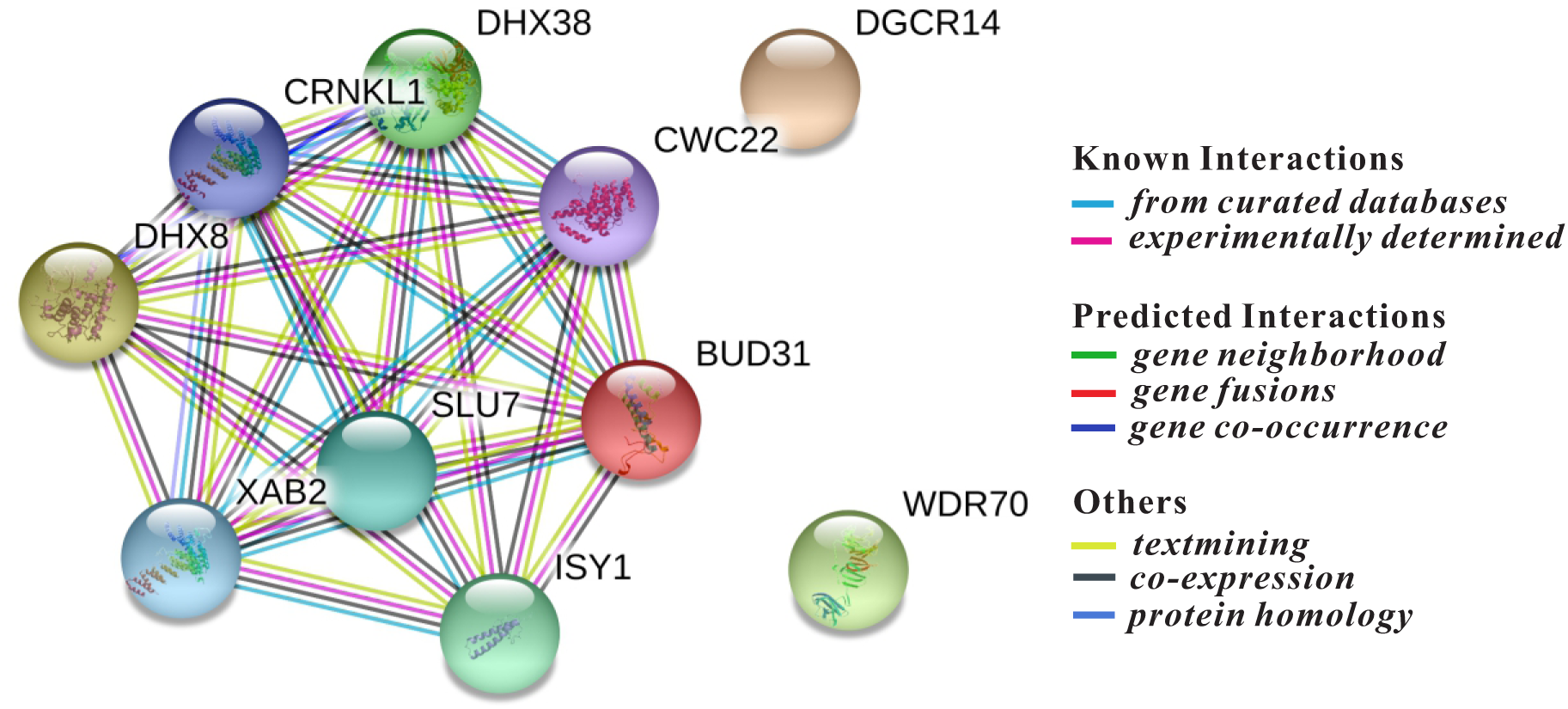
Protein interaction network of confirmed screening hits. The 10 validated hits were analyzed by the online database STRING for protein-protein interactions. The graph shows known and predicted protein-protein interactions for the most enriched pathway, the “mRNA splicing” pathway (pathway ID: GO 0000398; false discovery rate: 5.07e-14; minimum required interaction score: 0.400 (medium confidence)).

### CRINKL1 and other cellular factors impact on HIV-1 unspliced and fully-spliced mRNA levels

To follow-up on the reduction of splicing as the potential mechanism of enhanced Gag-BFP expression, the top 5 hits and WDR70 were analyzed for their effect on HIV-1 mRNA splicing. Attempts to generate J-dual#3 cells with a stable knock-out of the six selected target genes were not successful, presumably due to the fact that the selected genes are essential for cell viability (53). We therefore analyzed HIV RNA splicing and nuclear export after knock-down of these genes in short-term cultures. J-dual#3 cells were transduced with the lentiviral vectors encoding the different guide RNA genes and Gag-BFP positive cells were sorted to enrich for cells with the desired gene knock-down. Total RNA was then extracted from the sorted cells and the copy numbers of unspliced HIV RNA per ng extracted total RNA were determined and compared to the copy numbers of unspliced RNA in J-dual#3 cells transduced with a negative control vector (NT1). Consistent with the stronger BFP expression, unspliced HIV RNA copy numbers were also enhanced 3- to 9-fold (Fig. 4A). Fully-spliced HIV-1 RNAs were also quantified, revealing a marginal decrease in cells transduced with XAB2 and WDR70 targeting sgRNAs, but not with the other gene targeting sgRNA (Fig. 4A). Differential effects of the gene targeting on HIV-1 FS and HIV-1 US RNA also lead to decreased ratios of FS over US transcripts (Fig. 4A). For comparative reasons we also transduced J-dual#3 cells in an independent experiment with a lentiviral vector expressing Rev, sorted Gag-BFP positive cells and quantified HIV-1 US and FS transcripts. Rev enhanced the US HIV-1 RNA levels approximately 5-fold, which coincided with a minor reduction in FS transcripts (Fig. 4A).

**Figure 4.**
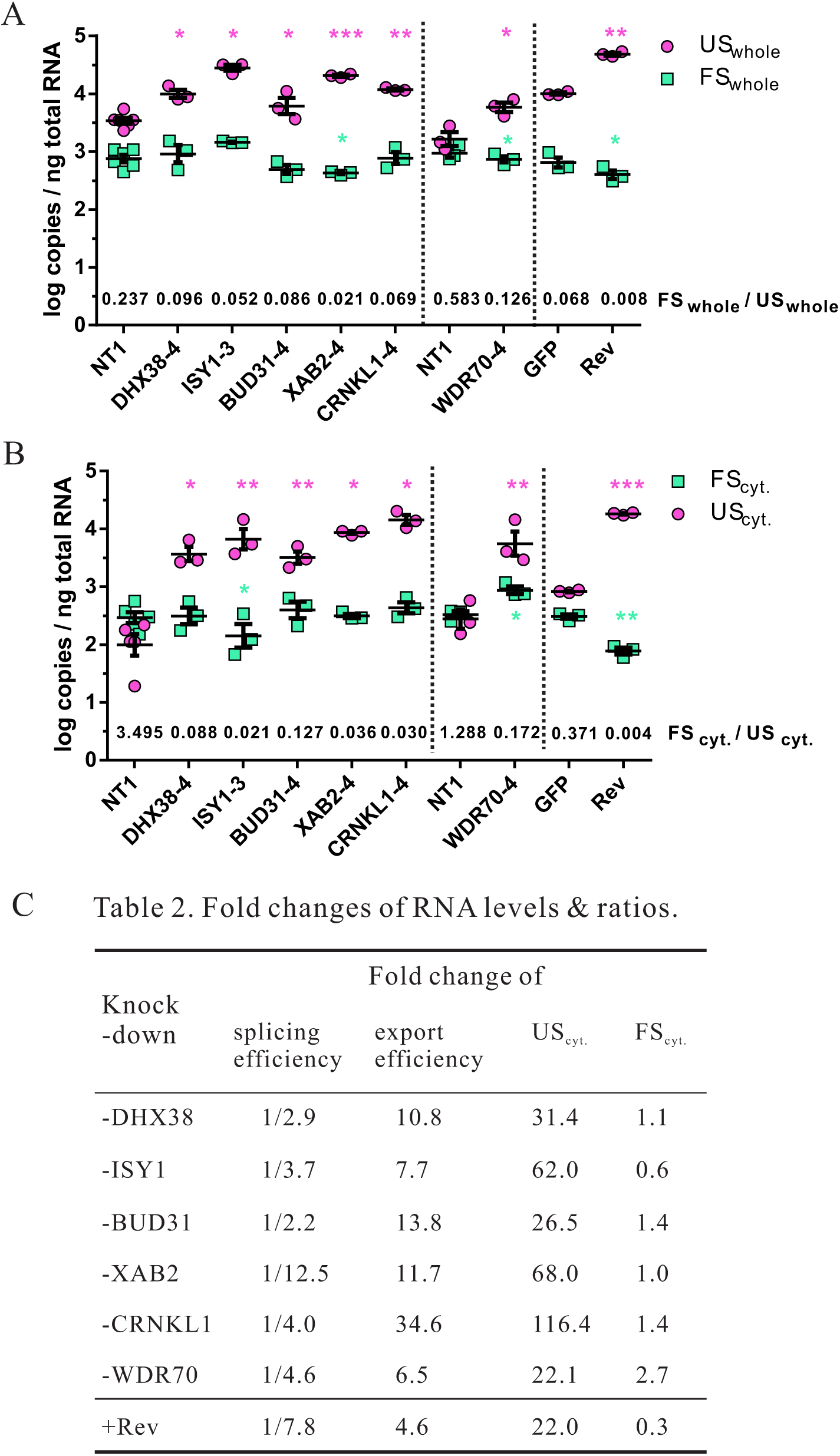
Role of confirmed screening hits in HIV-1 RNA splicing and export. (A) J-dual#3 cells were co-infected at an MOI of 3 with a lentiviral vector encoding Cas9 and a lentiviral vector expressing the guide RNAs indicated. In addition, J-dual#3 cells were transduced by a lentiviral vector encoding GFP or Rev at an MOI of 1. At 4 days after infection, BFP positive cells were sorted by flow cytometry from cells transduced with lentiviral vectors encoding Rev or the guide RNAs targeting the confirmed hits. Matched controls expressing NT1 or GFP were used without sorting. Total RNA was purified from whole cell extracts. HIV-1 US and FS RNAs copy numbers in these samples were quantified by RT-qPCR and are shown as log copies per ng total RNA extracted from whole cells. Log copies from at least three experiments, as well as their mean and SEM are shown. The mean of the ratio of FS_whole_/US_whole_ of each experiment is shown above the x-axis for each of the treatment groups. (B) J-dual#3 knockout cells and control cells were prepared as described in (A), but total RNA was extracted from the cytoplasmic fraction of each cell sample. Log copies per ng total cytoplasmic RNA and mean ratios of FS_cyt._/US_cyt._ were calculated and presented as in (A). (C) The fold changes in the different treatment groups for the splicing efficacy (FS_whole_/US_whole_), the nuclear export efficacy (US_cyt._/US_whole_), and cytoplasmic levels of unspliced (US _cyt._) and fully spliced (FS _cyt._) transcriptes were calculated by dividing the values in the treatment groups by the ones obtained from the matched control cells (NT1, GFP) as described in the Materials and Methods. Paired t-test was performed on the log copy numbers/ng total RNA between each of the treatment groups and the matched control. * p<0.05, ** p<0.01, ***p<0.001. Unless otherwise indicated, differences were not significant. Dotted lines separate experiments that were not performed in parallel and therefore contain independent controls. WDR70.KD cells were sorted on day 7 after transduction, while other cells were sorted on day 4.

The splicing process is closely linked to nuclear mRNA trafficking (54, 55). Therefore, we also determined the effect of targeting the six selected candidate genes on the cytoplasmic levels of HIV-1 US and FS RNA (Fig. 4B). Proper separation of the cytoplasm from nuclear components was controlled by examining GAPDH pre-mRNA levels and a nuclear marker protein in the cytoplasmic fraction from Jurkat cells (56, 57). GAPDH pre-mRNA levels per ng total RNA extracted from the cytoplasm were approximately 50-fold lower than GAPDH pre-mRNA levels per ng total RNA extracted from whole cells. Consistently, the nuclear LaminB protein could not be detected in the cytoplasmic fractions (Fig. S4).

After transduction with the different targeting vectors, HIV-1 US RNA in the cytoplasm of Gag-BFP positive cells was enhanced approximately 22 to 116-fold, while the cytoplasmic FS RNA levels were affected only 0.6 to 2.7-fold. The magnitude of the enhancement of cytoplasmic levels of HIV-1 US RNA exceeded the enhancement of HIV-1 US RNA in whole cell extracts, indicating that knocking-down the selected target genes not only reduced splicing but also enhanced nuclear export of the HIV-1 US RNA. Strong enhancement of cytoplasmic HIV-1-US RNA export was also observed in Rev-expressing cells. However, in contrast to the cells transduced with the targeting vectors, cytoplasmic FS RNA levels in Rev-transduced cells were significantly reduced, consistent with the previously reported suppression of the TAP/NXF1 mediated RNA export by Rev (58).

To dissect the relative contributions of inhibition of splicing and enhancement of nuclear export of HIV-1 US RNA, we compared the ratio of HIV-1 FS RNA to HIV-1 US RNA in whole cell extracts as a measure of splicing efficiency and the ratio of HIV-1 US RNA in cytoplasmic extracts to the HIV-1 US RNA in whole cell extracts as a measure of nuclear export efficiency. This revealed two different patterns of responses. The fold-reduction of the splicing efficiencies in cells transduced with ISY1, XAB2 and WDR70 targeting vectors mirrored the fold-enhancement of the respective nuclear export efficiencies. Expression of Rev induced a similar pattern of response. In contrast, transduction with DHX38, BUD31 or CRNKL1 targeting vectors predominantly enhanced the nuclear export efficiency with at least three-fold lower effects on splicing efficiency (Fig. 4C). In particular, knock-down of CRNKL1 enhanced the ratio of HIV-1 US RNA in cytoplasmic extracts to HIV-1 US RNA in total cell extracts more than 34-fold, while the ratio of HIV-1 FS RNA to HIV-1 US RNA in whole cell extracts was only decreased 4-fold. Targeting of CRNKL1 also led to the strongest enhancement of cytoplasmic levels of HIV-1 US RNAs without reducing cytoplasmic HIV-1 FS RNA levels (Fig. 4C), indicating that CRNKL1 is a major nuclear retention factor of the genomic HIV-1 US RNA.

### CRNKL1 knock-down shifts cytoplasmic levels HIV-1 splice variants

To confirm the role of CRNKL1 in the regulation of cytoplasmic HIV RNA levels and to explore potential effects of CRNKL1 on cytoplasmic levels of cellular RNAs a transcriptomic analysis was performed after a short-term knock-down of CRNKL1. J-dual#3 reporter cells were again transduced with the CRNKL1 targeting vector and Gag-BFP-positive cells were sorted by flow cytometry. RNA was extracted from the cytoplasmic fraction of sorted cells and J-dual#3 cells transduced with a control vector. The extracted RNAs from 4 biological replicates each were then enriched for mRNAs and sequenced by the Illumina High-seq procedure. In addition to this, cytoplasmic mRNA was extracted from J-dual#3 cells expressing Gag-BFP after transduction with the Rev-encoding lentiviral vector and sequenced.

To confirm the effectiveness of Cas9/sgRNA-mediated knock-down by transcriptomic analysis, a core target region in exon 3 of the CRNKL1 gene was defined as nucleotides −3 to +3 relative to the SpCas9 cleavage site. The number of RNA-seq reads that contained 10 nucleotides upstream and downstream of the core target region were then determined (Fig. S5). While 55 reads on average mapped to the region flanking the core target region in the control cells, only a mean of 5.8 reads were detected in cells transduced with the CRNKL1 knock-down vector (Fig. S5). Larger deletions and substitutions outside the core target region probably reduce the number of reads that can be mapped to the flanking region. Comparison of the reads from CRNKL1 knock-down cells that can be mapped to the flanking region with the wild type CRNKL1 core target region revealed insertions, deletions and point mutations in 22 out of 23 reads. In control cells, only 2 out of 219 reads contained single point mutations clearly indicating efficient inhibition of CRNKL1 expression after Cas9/sgRNA targeting (Fig. S5).

HIV reads were mapped to the proviral DNA of HIV-dual-GT revealing more reads mapping to intronic regions after CRNKL1 knock-down and Rev expression than in the controls (Fig. 5A). The percentage of reads mapping to HIV among all the reads uniquely mapping to either the human or the viral genome (total uniquely mapped reads) was enhanced two fold in CRNKL1 knock-down cells (Fig. 5B). This was not due to an enhanced transcriptional activity since the percentage of reads mapping to Exon 1, which is shared by all HIV transcripts, was only affected marginally(Fig. 5C). Changes in HIV-1 US RNA were assessed by determining the percentage of reads mapping to the first intron of HIV-1 between SD1–SA1 (5D) or mapping to the unspliced SD1 site (5E) revealing 4 and 12-fold increases under CRNKL1 knock-down conditions, respectively. The percentage of reads mapping to the unspliced SD4 site and therefore containing the *env* intron also increased by a factor of 9 (Fig. 5F). The change in the percentage of all spliced HIV reads was negligible (Fig. 5G) indicating that knock-down of CRNKL1 does not reduce splicing of HIV RNAs in general.

**Figure 5.**
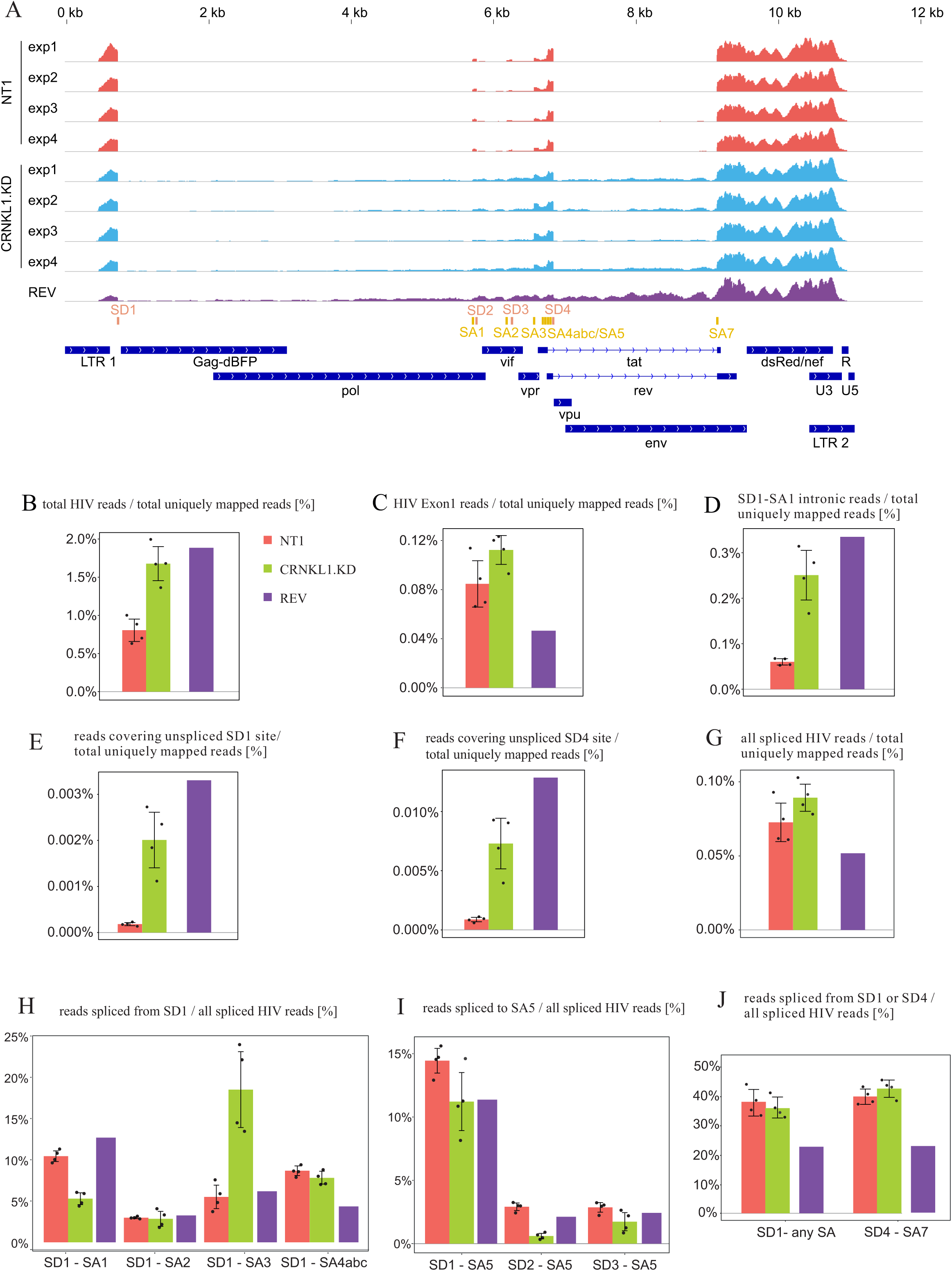
Effect of CRNKL1 depletion on HIV RNA splicing efficiency, accuracy, and export. (A) Alignments of reads from the transcriptomic analysis to HIV-dual-GT. Shown are coverage profiles for the individual RNA-seq samples, colored, from top to bottom the non-targeting control (NT1, red), CRNKL1 knock-down (cyan), and rev overexpression (purple). Splice sites are shown in orange (donors) and dark yellow (acceptors). Coding regions and LTRs are indicated in dark blue. (B) Percentage of all HIV reads among the total (HIV + human) uniquely mapped reads. (C) Percentage of reads mapping to HIV Exon 1 (from the transcription start site to SD1) among the total uniquely mapped reads. (D) Percentage of reads in the first intron between SD1 and SA1 among the total uniquely mapped reads. (E) Percentage of reads across the unspliced SD1 site among the total uniquely mapped reads. (F) Percentage of reads across the unspliced SD4 site among the total uniquely mapped reads. (G) Percentage of reads spanning SD and SA sites (spliced reads) among the total uniquely mapped reads. (H) Percentages of reads spanning SD1 and the indicated splice acceptor sites among all spliced HIV reads. For SD1-SA4abc, reads spliced from SD1 to SA4a, SA4b and SA4c are summed up. (I) Percentages of reads spanning the indicated splice donor sites and SA5 among all spliced HIV reads. (J) Percentage of reads spliced from SD1 to any SA site or from SD4 to SA7 among all spliced HIV reads. (B-J) Shown are averages and standard deviations for the control and CRNKL1 knock-down samples. Individual values are shown as black dots. For the rev overexpression sample (purple), the individual value is shown.

To further explore differences in cytoplasmic levels of differentially spliced HIV-1 transcripts the precise splice site usage was also systematically analyzed by calculating the percentage of reads covering specific SD-SA pairs among all spliced HIV reads (Fig. S6A,B). This revealed that knock-down of CRNKL1 reduced cytoplasmic HIV transcripts spliced from SD1 to SA1 approximately two-fold, while transcripts spliced from SD1 to SA3 were enhanced three-fold (Fig. 5H). To further assess the effect of the CRNKL1 knock-down on the major Env encoding transcript, the percentage of reads spliced to SA5 was determined (Fig. 5I). Independent of the SD sites, usage of SA5 trended to be reduced, which is similar to an effect recently reported for the splicing factor DDX17 (59). Moreover, the level of fully spliced transcripts seemed unaffected by the CRNKL1 knock-down since the percentages of reads spliced either from SD1 to any splice acceptor site or from SD4 to SA7 were nearly the same (Fig. 5J).

### CRNKL1 is a regulator of cytoplasmic cellular mRNA levels

The influence of the CRNKL1 knock-down on cytoplasmic mRNA levels was also determined (Fig. S7A,B). To display the results the log2-transformed expression differences were plotted against the expression level for each gene in a MA-plot (Fig. 6A). To reduce noise and arbitrarily large fold changes from low expressed genes the log2 fold changes are shrunk putting less weight on low expressing genes. Significant differences after correction for multiple testing revealed the upregulation of 1249 and downregulation of 2569 mRNAs, respectively. Two hundred and ten mRNAs even showed a more than 4-fold (log2 fold change >2) increase after knock down of CRNKL1. Interestingly, upregulated mRNAs were enriched for the functional categories involved in transcription (Fig. 6B) while downregulated mRNAs were more frequently associated with functions in cell cycle and organelle organization (Fig. 6B).

**Figure 6.**
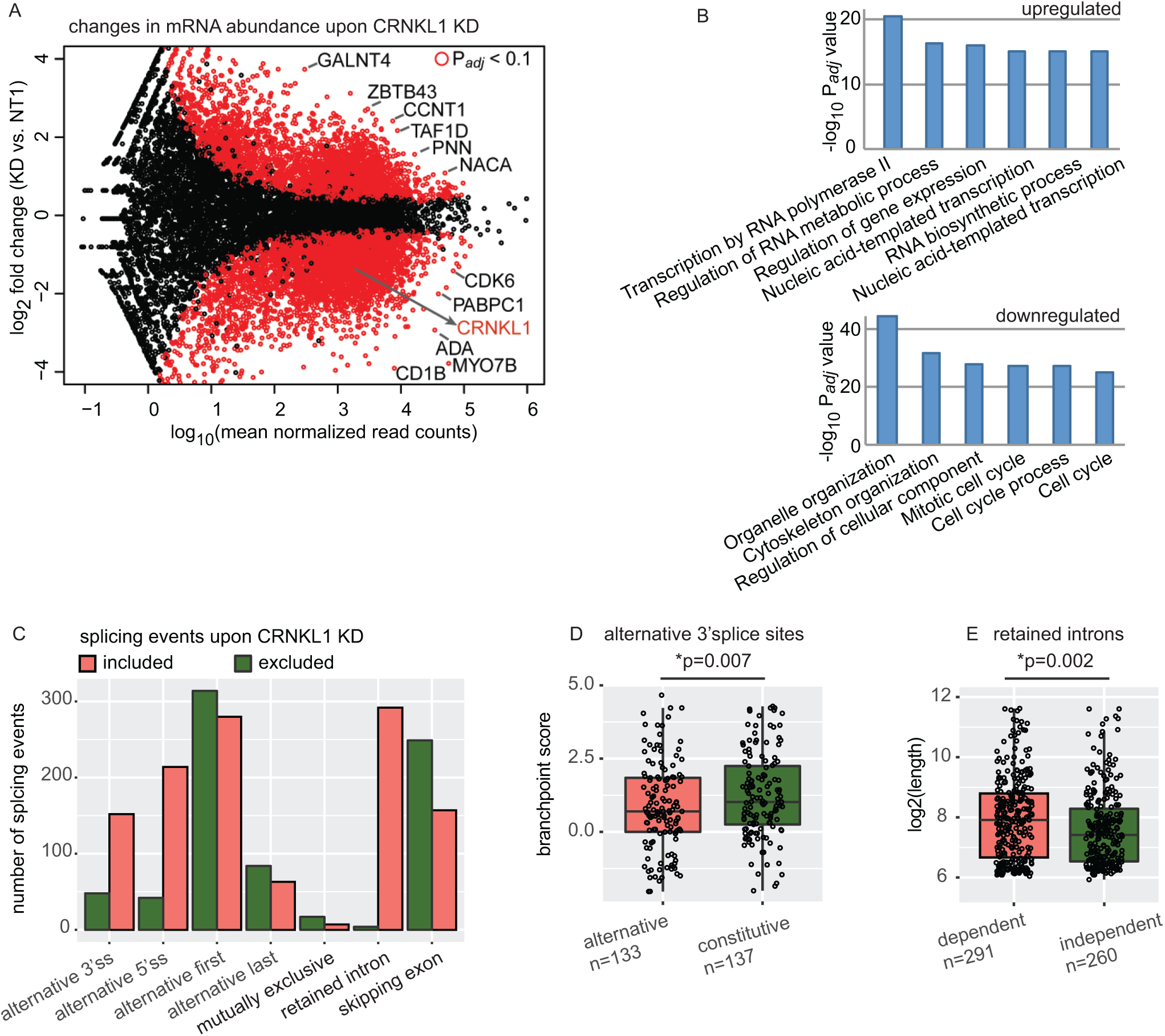
Changes in the host transcriptome upon CRNKL1 knock-down. (A) RNA-seq differential expression analysis of mRNA abundance upon CRNKL1 KD. MA plot shows differences in the abundance of protein-coding transcripts in CRNKL1 KD vs. control (NT1) conditions. mRNAs with significant differences are depicted in red (Padj <0.1). (B) Gene Ontology enrichment analysis of mRNAs with upregulated and downregulated expression upon CRNKL1 KD. Top enriched GO Biological Process terms are shown. (C) Upregulated (included) (ΔPSI (KD-NT1) >0.25, Padj <0.01) and downregulated (excluded) (ΔPSI (KD-NT1) <-0.25, Padj<0.01) splicing events were quantified from the RNA-seq CRNKL1 KD experiment. Bar plot shows absolute numbers of splicing events that are up or down regulated upon CRNKL1 knock-down. (D) Branch point scores were calculated by BPfinder and alternative 3’splice sites with higher usage in CRNKL1 KD condition were compared to 3’splice sites of the same genes with unchanged (constitutive) usage. Box plots and data points are shown. Wilcoxon rank sum test was used to test for significance. Absolute numbers of 3’ splice sites included in the analysis for each group are indicated. (E) Length of retained introns with higher inclusion upon KD (CRNKL1-dependent) was compared to the length of unchanged introns upon KD (CRNKL1-independent), which were located in the same transcripts as CRNKL1-dependent ones. Box plots and data points are shown. Wilcoxon rank sum test was used to test for significance. Absolute numbers of retained introns analyzed in both groups are indicated. PSI, percent spliced-in for splicing events.

Given the close link between RNA export and splicing, a more general role of CRNKL1 on cytoplasmic levels of splice variants was analyzed upon CRNKL1 knock-down. Individual splicing events (including skipped exons, alternative splice sites and retained introns) were quantified using the percent spliced-in metric (PSI, Schafer et al., 2015) and differential analysis was carried out by SUPPA (61). Of all quantified splicing events in cytoplasmic mRNAs upon CRNKL1 depletion, 2.4 % were upregulated and 1.6% downregulated (Fig. S7C,D). Introns were more frequently retained upon CRNKL1 depletion compared to wild-type cells (Fig. 6C). Similarly, alternative 3’ and 5’ splice sites were more frequently detected upon CRNKL1 knock-down, suggesting that CRNKL1 inhibits usage of alternative splice sites (Fig. 6C).

A total of 295 introns were more frequently retained in cytoplasmic mRNAs after CRNKL1 knock-down. Further analyses revealed that not all introns of the same transcript were affected in the same way (Fig. S8), but rather that intron retention was highly specific for one or a few introns of the same gene. Enhanced cytoplasmic levels of intron-retaining mRNAs could be confirmed by real-time PCR with primers spanning the respective exon – intron junctions (Fig. S8).

Genes containing introns or alternative splice sites that were upregulated after CRNKL1 knock-down were not enriched for particular functional categories (data not shown). Enhanced detection of introns or alternative splice sites in cytoplasmic mRNAs after CRNKL1 knock-down indicates that CRNKL1 suppresses either these splicing events or the nuclear export of the respective splice variants. To gain insight into the principles of regulation of these CRNKL1 knock-down dependent introns and alternative splice sites, we analyzed common features of these splicing events, including splice site strength, branch point and polypyrimidine tract score, GC content and intron length. CRNKL1 knock-down-dependent alternative 3’ splice sites showed significantly lower branch point scores than constitutive 3’ splice sites (Fig. 6D), but a similar length of polypyrimidine tracts (Fig. S7E). We also found that CRNKL1-dependent introns were significantly longer than CRNKL1-independent introns within the same transcript (Fig. 6E) and trended to have longer polypyrimidine tracts (Fig. S7F) but showed no difference in GC content (Fig. S7G).

## Discussion

Using a genome-wide screen for cellular factors repressing expression of an HIV-1 structural protein from the unspliced RNA, we identified ten target proteins linked to mRNA metabolism, with nine of them associated with the spliceosome. Interestingly, five of them (ISY1, BUD31, XAB2, CRNKL1 and CWC22) are found in the Prp19-associated complex. This complex is important for stabilizing the spliceosome, which is also involved in DNA repair, transcriptional elongation, and RNA export (51). Given that the yeast homolog of WDR70 was reported to interact with Prp19 (62) and is now shown to affect RNA splicing, it could be a new member of the Prp19-associated complex. Due to the close linkage of splicing with RNA stability and nuclear RNA export, it is difficult to disect the precise mechanism by which deletion of the identified nuclear proteins enhance cytoplasmic RNA levels of HIV-1-US RNA.

Theoretically, enhanced transcription, reduced nuclear or cytoplasmic degradation, decreased splicing, and/or enhanced nuclear export can increase cytoplasmic levels of HIV-1 US RNA (US_cyt._) (Fig. 7). Enhanced transcription and decreased nuclear degradation seem unlikely since the percentage of reads mapping to the first HIV-1 Exon present on all viral transcripts was not significantly enhanced (Fig. 5B). The absence of enhanced cytoplasmic levels of FS RNAs and DsRed expression levels from the dual reporter construct (Fig. 4B, Fig. S2A) also argues against such a mechanism. Assuming a single pool of nuclear HIV-1 US RNA that could follow either of the two remaining pathways, differential effects on the nuclear export efficiency (US_cyt._/US_whole_) and splicing efficacy (FS_whole_/US_whole_) are expected (Fig. 7). Enhancing nuclear export of HIV-1 US RNA should strongly enhance the ratio of US_cyt._/US_whole_. Since rapid export may reduce splicing and degradation, the ratio of FS_whole_/US_whole_ may responsively decrease, although to a lower extent. Reduced splicing should primarily decrease the ratio of FS_whole_/US_whole_, but not the ratio of US_cyt._/US_whole_ (Fig. 7). Similar to an enhanced nuclear export, reduced degradation of cytoplasmic HIV-1 US RNA should also lead to enhanced US_cyt._/US_whole_ and decreased FS_whole_/US_whole_ ratios (Fig. 7), but the association of the identified proteins with the spliceosome localized in the nucleus argues against such a cytoplasmic effector mechanism.

**Figure 7.**
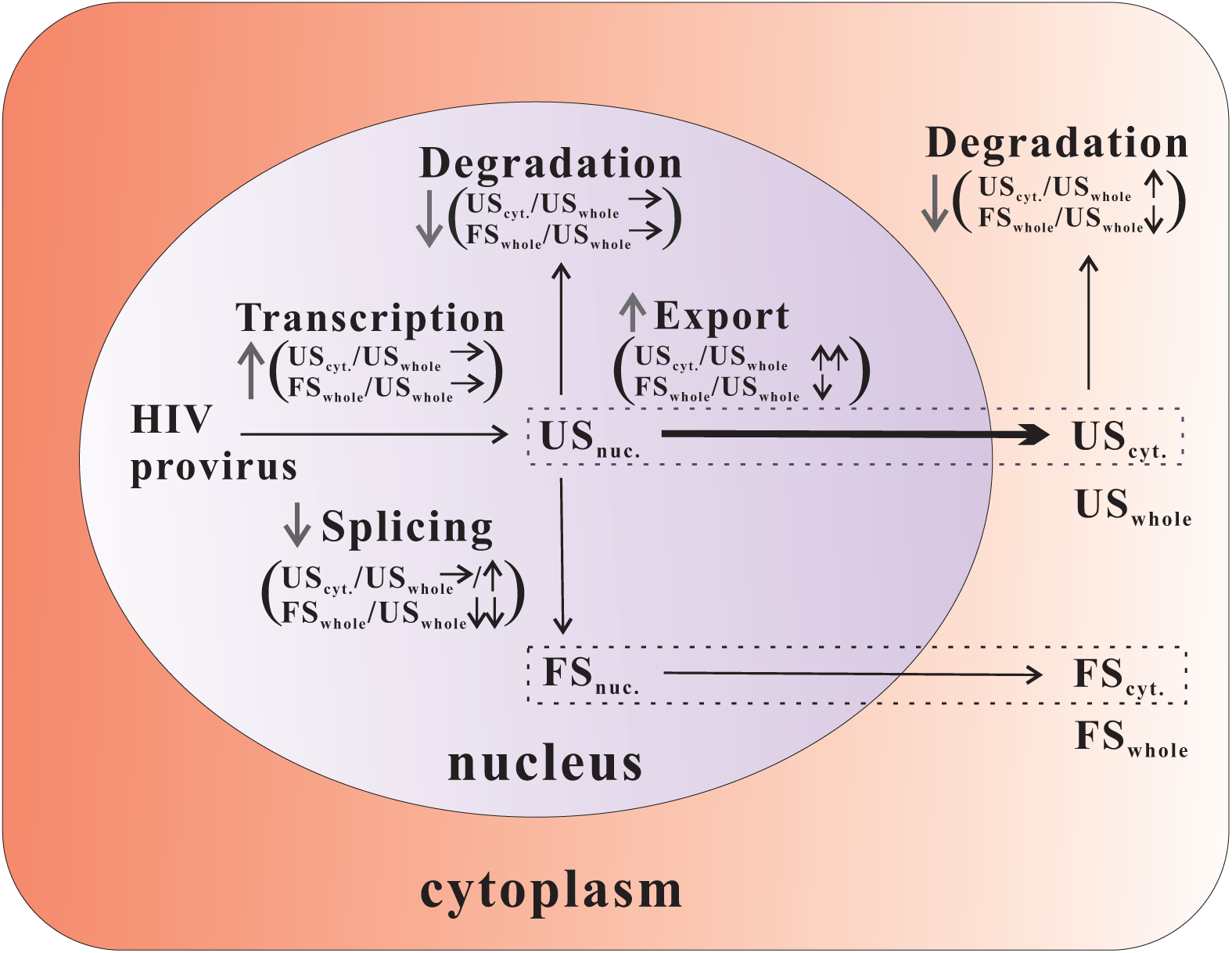
Potential mechanisms leading to increased cytoplasmic levels of unspliced HIV-1 RNA. Graphical representation of HIV-1 RNA metabolism and expected consequences of up-(↑) or down-(↓) regulation or no change (→) of the respective steps on the nuclear export efficacy (US_cyt._/US_whole_) and splicing efficacy (FS_whole_/US_whole_). The nuclear export pathway is highlighted in bold.

Clearly, ablation of all 6 target genes tested reduced the splicing efficacy (FS_whole_/US_whole_) indicating that the targeted genes are enrolled in splicing and/or nuclear retention. The relative contribution of reduced splicing versus enhanced nuclear export of the HIV-1 US RNA differed between the different target genes. The XAB2 knock-down decreased the splicing efficacy 12.5-fold and the nuclear export efficacy of the HIV-1 US RNA 11.7-fold. Since splicing and nuclear export efficacy were affected to a similar degree, it is not possible to conclude whether XAB2 predominantly acts as a splicing or nuclear retention factor. This is consistent with previous observations that depleting splicing factors (e.g. SF1, U2AF65 and Prp19) can cause pre-mRNA leakage into the cytoplasm (30, 63–65). In contrast to the effect of the XAB2 knock-down, the splicing efficacy after knock-down of CRNKL1 is only decreased 4.0-fold, while the nuclear export efficiency based of the HIV-1 US RNA increases 34.6-fold. Since the ratio of HIV-1 FS to US transcripts in the cytoplasmic RNA is also reduced to a larger extent than in the whole cell RNA, enhanced nuclear export of the HIV US RNA after inactivation of the target gene seems to play a dominant role. This indicates that CRNKL1 is primarily required for retention of the HIV-1 US RNA in the nucleus and not for its splicing. The precise molecular mechanism by which the CRNKL1 knock-down leads to a more than 100-fold enhancement of cytoplasmic HIV US RNA levels remains to be clarified. CRNKL1 has been previously reported to regulate splicing and assist the export of cellular intron-less mRNAs in cooperation with the TREX (TRanscription-EXport) complex, U2AF65 and other NTC members (66–68). Our observation that targeting of CRNKL1 did not enhance cytoplasmic levels of HIV-1 FS RNA, which uses the default TAP/NXF1 export pathway, indicates that CRNKL1 is not a global nuclear mRNA retention factor. Our transcriptomic analysis also indicates a highly selective differential regulation of cytoplasmic levels of HIV-1 splice variants. The only spliced HIV-1 transcripts that were clearly upregulated by CRNKL1 knock-down have spliced SD1 to the SA3 site (Fig. 5G) and may encode Tat. Whether this represents an upregulation of *env* intron retaining transcripts or the tat-encoding fully-spliced transcript is unclear. The enhanced percentage of cytoplasmic HIV transcripts harbouring an unspliced SD4 site (Fig. 5F) and the unchanged percentage of transcripts harbouring a spliced SD4 site (Fig. 5J) argue for the former.

The transcriptomic analyses after CRNKL1 knock-down also revealed a selective regulation of cytoplasmic levels of cellular mRNAs. We found that 11.0% and 20.1% of expressed mRNAs in the cytoplasm were up- and downregulated, respectively. Functionally, the upregulated mRNAs are associated with transcription, while downregulated mRNAs are more frequently associated with the cell cycle and organelle organization suggesting that CRNKL1 participates as a master regulator in these cellular programs. Since altered nuclear export after CRNKL1 depletion could result in differences in the abundance of different cytoplasmic mRNA isoforms, we investigated alternatively spliced cytoplasmic isoforms upon CRNKL1 knock-down. Upon CRNKL1 depletion a much higher number of included introns and alternative 3’ and 5’ splice sites than the excluded ones were detected, suggesting that CRNKL1 promotes splicing of these events or highly selective nuclear retention. Intron retention in the absence of CRNKL1 was highly restricted and even varied among introns of the same gene (Fig. S8). Searching for common properties of introns that were retained revealed that their mean length was significantly greater compared to introns with unchanged inclusion upon CRNKL1 depletion. In addition, usage of 3’ splice sites with lower branch point scores seems to be blocked in the presence of CRNKL1. Therefore, the sequence features of CRNKL1-dependent splicing events seem to reside at the 3’ rather than at the 5’ end of the intron.

Mechanistically, one plausible hypothesis based on the nuclear retention observed for the HIV-1 unspliced RNA is that a nuclear RNP complex binds in a CRNKL1 dependent manner to RNA motifs present in a highly selective subset of cellular and viral introns leading to their nuclear retention and splicing. CRNKL1 dependent binding to binding motives on cellular transcripts leading to nuclear retention and degradation may also explain the increase in cytoplasmic levels of cellular mRNAs upon CRNKL1 knock-down.

In summary, our study identifies three spliceosomal proteins that are required for nuclear retention of the HIV-1 unspliced RNA and indicates that CRNKL1-dependent nuclear retention is a novel mechanism for the regulation of cytoplasmic levels of intron-retaining cellular mRNAs that is hijacked by HIV-1.

## Materials and Methods

### Plasmids

Plasmids expressing viral proteins, HIV-1 Rev (pcRev), HIV-1 Tat (pcTat), HIV-1 codon-optimized Gag-Pol (Hgp^syn^) and VSV-G (pHiT-G) were described previously (23, 56). Lentiviral packaging plasmid pPAX2 and pMD2.G were obtained from Addgene (#12260 and #12259). HIV-1 proviral plasmid HIVΔe is based on the HIV-1 molecular clone NL4-3 and contains a 4-nucleotide deletion at its NheI site leading to inactivation of *env*. The plasmid HIVΔr-e contains an additional mutation of the start codon of *rev* and a premature stop codon in the first *rev* exon (57).

Proviral reporter plasmid HIV-dual-GT was constructed based on HIVΔr-e. A cassette containing mTagBFP2, a PEST coding sequence (destabilized BFP) and two stop codons were ligated in frame to the 3’ end of *gag*, forming a new ORF for Gag-BFP. The Slippery Sequence TTTTTTA (nt 2085 to nt 2091 of the GenBank entry AF324493.2) was mutated to CTTCCTG to prevent Gag-Pol translation (69). The coding sequence of Nef close to its 5’end (nt 8796 to nt 8880 of the GenBank entry AF324493.2) was replaced by a cassette consisting of a 3 x Ala linker sequence and the DsRed max coding region followed by two stop codons. The sequence (nt 5131 to nt 6297 of the GenBank entry AF324493.2) ranging from *vif* to *vpu* was replaced by the corresponding region of NL4-3-Δ6-drEGFP (70) harboring inactivating point mutations in *vif*, *vp*r and *vpu*. The *rev* ORF was inactivated as described above for HIVΔr-e by PCR mutagenesis. The plasmid HIV-dual-GT including the final sequence is available from AddGene (plasmid ID 122696). Individual CRISPR knock-out plasmids were constructed based on lentiCRISPRv2 (#52961, Addgene) and GeCKOv2 sgRNA target sequences. All sgRNA sequences of the GeCKO v2 library can be found at www.addgene.org/pooled-library/zhang-human-gecko-v2/. The different sgRNA sequences selected for validation are provided in Supplementary Table S1A. Each of the sgRNA sequences were cloned into the vector following the instructions on the Addgene webpage (www.addgene.org/52961/).

### GeCKO v2 library

The pooled Human GeCKO v2 CRISPR knockout plasmid library was a gift from Feng Zhang (41) and obtained from Addgene (#1000000048 and #1000000049). It is supplied as two sub-libraries, each containing 3 different sgRNA constructs targeting each gene in the genome. Amplification of the lentiviral vector plasmid library was performed essentially as described by the contributors’ protocol (41). The complexity of the library was confirmed by HiSeq NGS of the pooled plasmid library (data not shown).

### Cell culture

HEK293T (DSMZ) and TZM-bl (71) (NIH AIDS reagent program) cell lines were maintained in DMEM medium supplemented with 10% FCS and 1% penicillin/streptomycin. Jurkat, Clone E61 (ATCC) and reporter cell lines J-dual#3 and J-dual#6 were maintained in RPMI 1640 supplemented with 10% FCS, 1% penicillin/streptomycin. After expansion, J-dual#3 and J-dual#6 stocks were stored frozen in aliquots at −80°C. A fresh aliquot was thawed and expanded in each assay.

### Cell transfection

HEK293T cells were routinely transfected by the polyethylenimine (PEI)-precipitation method (56). In general, 3.5 million cells per T25-flask (CELLSTAR) were seeded 24 h before transfection. For each transfection, plasmid DNA was mixed with calf thymus carrier DNA (Thermo Fisher Scientific) to give a total of 10 μg DNA. The 10 μg DNA were mixed with 15 µl PEI (1µg/µl, Sigma Aldrich) and added to the cells. After 8 h of incubation with the cells, the transfection medium was exchanged with fresh culture medium. Supernatants or cells were harvested 48 h post transfection. To produce pseudotyped HIVΔe, HIVΔr-e and HIV-dual-GT vector particles, 3 μg proviral plasmid, 3 μg Hgp^syn^, 2 μg pHiT-G, 1 μg pcTat and 1 μg pcRev were used. To produce individual CRISPR-KO lentiviral vector particles, 5 μg lentiCRISPRv2 plasmid, 3.75 μg pPAX2 and 1.25μg pMD2.G were used. To produce the pooled human GeCKO v2 lentiviral vector library, 24 millions of 293T cells were seeded into a T175 tissue culture flask. 24 h later, 20 μg GeCKO v2 library plasmids, 15 μg pPAX2, 5 μg pMD2.G, and 120 μl GenJet transfection reagent (SignaGen) were used for transfection. HIV-dual-GT-transduced Jurkat cells were transfected by the electroporation method described elsewhere (72). In brief, 5.0 millions of stably transduced cells were electroporated with 50 μg plasmids by the Gene Pulser X cell^TM^ Electroporation System (Bio-Rad) at 250 V and 1500 microfarads. 48 h post transfection, cells were harvested for flow cytometry analysis. The transfection efficiency after expression of a GFP reporter plasmid was estimated to be between 10% - 20%.

### Western blot analyses

Immunoblotting for HIV-1 Gag/CA p24 was described previously (57). Immunoblotting for Rev was performed with primary antibody Sheep Anti-Rev (1:4,500, #H6006, US Biological Life Sciences) and secondary antibody Rabbit Anti-Goat Ig/HRP (1:5,000, #P0160, Dako). Immunoblotting for α-Tubulin was performed using polyclonal Rabbit Anti-α-Tubulin antiserum as primary antibody (1:4,000, #600-401-880, Rockland), and Swine Anti-Rabbit Ig/HRP (1:5,000, #P0217, Dako) as secondary antibody. In the experiment of Figure 1B, anti-α-Tubulin blotting was conducted after stripping the blotted anti-CA membrane by stripping buffer (#2504, Millipore). The membrane was then re-blocked, and subsequently re-blotted with antibodies as described above. Immunoblotting for LaminB was performed with primary antibody Goat anti-LaminB (C-20) (1:500, #sc-6216, Santa Cruz Biotechnology) and secondary antibody Rabbit anti-goat Ig HRP (1:5,000, #P0160, Dako).

### Lentiviral vector preparation, infection and titration

Lentiviral vector particles were produced by transfection of 293T cells with the plasmids mentioned above. HIV-1 proviruses were packaged with a 3^rd^ generation packaging system (Hgp^syn^, pcRev, pcTat and pHiT-G) (23, 73, 74). CRISPR-KO constructs and a pool of the two GeCKO v2 plasmid A and B sublibraries were packaged with a 2^nd^ generation packaging system (pPAX2 and pMD2.G) (42, 75) for higher infectious titers. 48 h after transfection, the supernatants from transfected 293T cells were centrifuged at 1,500 rpm for 5 min to remove cell debris, and afterwards filtered through 0.45 μm filters. Viral stocks were aliquoted and stored under −80°C. Two batches of the GeCKO v2 lentiviral vector library were prepared. HiSeq NGS analyses of sgRNA representation in cells transduced with both batches showed an almost 100% identification of the sgRNAs encoded by the two GeCKO v2 plasmid sublibraries.

Jurkat, J-dual#3 and J-dual#6 cells were infected with the lentiviral vectors or the lentiviral vector library by spinoculation. 1.5 million of cells were centrifuged with 1 ml vector preparation, in the presence of 8 μg/ml polybrene (Sigma-Aldrich), at 2,000 rpm, for 3 h, at 33°C. After spinoculation, the supernatants were removed and the cells were continued to be cultured.

VSV-G-pseudotyped HIVΔe, HIVΔr-e and HIV-dual-GT vector particles were titrated on TZM-bl cells as previously described (57). The titers were 5.8×10^6^ and 5.1×10^6^ TU/ml for two independent titrations of HIVΔe, 4.3×10^6^ and 5.7×10^6^ TU/ml for two independent titrations of HIVΔr-e and 6.4×10^5^and 5.0×10^5^ TU/ml for two independent titrations of HIV-dual-GT. VSV-G-pseudotyped HIV-dual-GT vector particles were also titrated on Jurkat cells. The lentiviral reporter vector was serially diluted to 1:1, 1:2, 1:4, 1:8 and 1:16. The infection was done by spinoculation method. At 48 hpi, cells were analyzed by flow cytometry to determine the percentage of DsRed positive cells in each dilution. Vector titers on Jurkat cells were calculated as follows for all dilutions: (initial cell number) X (percentage of DsRed^+^ cells) X (dilution factor). The highest value resulting from these dilutions was taken as vector titer on Jurkat cells. The determined titers for two independent titrations of HIV-dual-GT were 4.9×10^5^ and 6.9×10^5^ TU/ml. The individual CRISPR-KO vectors, as well as the GeCKO v2 lentiviral vector library, were titrated on J-dual reporter cells and parental Jurkat cells. The cells infected by the serially diluted virus were grown for 24 h, afterwards divided into two equal aliquots. One aliquot was then cultured in 2 µg/ml puromycin, the other without puromycin. After 30 h of growth, viable cells were counted by trypan blue staining. The transduction percentage was calculated in the following way: (number of living cells in treated population) x100/ (number of living cells in untreated population). For the GeCKO v2 lentiviral vector library the transduction percentage was multiplied by the number of cells exposed to calculate the titer for the different dilutions of the library. The highest value obtained from the different dilutions of the same batch of the lentiviral vector library was taken as its titer. The titers were 1.3×10^7^ TU/ml for the first batch and 2.2×10^6^ TU/ml for the second batch.

### Flow cytometry analysis and Cell Sorting

Flow cytometry analyses were done using the BD™ LSRII flow cytometer (BD Biosciences). FACS data were analyzed by the Flowjo software. Cell sorting was done on a MoFlo XDP (Beckman Coulter) at the FACS core facility of the medical faculty.

### Next-Generation Sequencing of sgRNA libraries

Genomic DNA was extracted from the sorted or the control cells either using the Quick-DNA Mini Kit or the Midi Plus Kit (Zymo Research). The integrated LentiCRISPRv2 vector sgRNA sequences were amplified through three rounds of PCR using the Taq DNA Polymerase S (high specificity) (Genaxxon Bioscience). All obtained genomic DNA was used as template in parallel reactions in PCR1, with the cycling conditions 94°C 2 min, then 18 cycles’ 95-55-70°C, 15-20-45 s, final extension 70°C for 3 min. PCR1 reactions of respective cell samples were then combined and 10 μl of the pooled PCR1 product was used as template in PCR2, consisting of 95°C 2 min, then 15 cycles’ 95-60-72°C, 15-20-30 s, final extension 72°C for 3 min. In the indexing PCR3, 10 ng of purified PCR2 product were subjected to 95°C 2 min, then 8 cycles’ 95-59-72°C, 15-20-30 s, final extension 72°C for 3 min. The primer sequences and primer combinations used for each sample at each step are presented in Tables S1B,C. After PCR2 and PCR3, PCR products were purified by the AMPure XP PCR-Cleanup Kit (Beckman Coulter); DNA concentrations were measured by the Qubit Quant-iT dsDNA HS Assay Kit (Thermo Fischer Scientific). Manipulations followed the manufacturers’ instructions. DNA libraries were sequenced on Illumina MiSeq (for enriched samples) or Ilumina HiSeq 2500 (for unselected control) instruments. NGS readouts were subjected to MAGeCK analysis (43).

### Cell fractionation, RNA isolation and RNA quantification

The cell fractionation protocol has been described previously (76). The plasma membrane was lysed with chilled NP40-Buffer consisting of 10 mM Hepes (pH 7.8 adjusted by KOH); 10 mM KCl; 20% Glycerol; 0.25% NP40. DTT was added to a final concentration of 1 mM just before use. After incubation on ice for a maximum of two minutes, the supernatant consisting of the cytoplasmic fractions was collected after centrifugation at 400 g, for 5 min, at 4°C. Cytoplasmic RNA was extracted by the TRIzol (Thermo Fisher Scientific) -Chloroform (Sigma-Aldrich) method (77) or by the Direct-zol RNA miniprep plus kit (Zymo Research). RNA from the whole cell extracts was purified using the RNA-Easy Kit (Qiagen). RNA extracted by TRIzol-Chloroform or the RNA-Easy kit was treated with DNaseI (NEB) to remove DNA contamination. Under the protection of 5 mM EDTA, DNaseI was inactivated at 75°C for 10 min. For the Direct-zol RNA miniprep plus kit purified RNA, DNA contamination was removed by in-column DNA digestion reagents supplied. Total RNA concentration was measured by the Qubit Quant-iT RNA HS Assay Kit (Thermo Fisher Scientific) according to the manufacturer’s instructions.

Reverse-transcription PCRs (RT-qPCR) for quantification of HIV-1 US, FS RNAs and cellular GAPDH pre-mRNAs were done with the QuantiTect SYBR Green RT-PCR Kit (Qiagen), and performed essentially as described previously (56, 78). Using the applied Biosystems 7500 Real-Time PCR machine cycling conditions were as follows: HIV-1 US RNA: 95°C 10 sec for denaturation, 65°C 60 sec for annealing, 72°C 30 sec for elongation and fluorescence detection; for HIV-1 FS RNA: 95°C 10 sec for denaturation, 64°C 15 sec for annealing, 72°C 15 sec for elongation, and 81°C 30 sec for fluorescence detection.

RT-qPCRs for relative quantification of Intron 6-retaining RPL10 mRNAs and Intron 1-retaining C19orf53 mRNAs were done with primers illustrated in Table S1D. Cycling conditions for both: 95°C 10 sec for denaturation, 50°C 60 sec for annealing, 72°C 60 sec for elongation and fluorescence detection.

### Calculation of the ratios of HIV FS and US RNA levels and fold changes

To measure the effect of candidate genes and Rev on RNA splicing efficacy, we first determined the ratio between FS_whole_:US_whole_ by dividing the FS copy numbers/ng extracted RNA by the US copy numbers/ng extracted RNA for each sample of each treatment group (knock-down or +Rev) and then calculated the mean of the ratios of replicates of each treatment group. The mean of the ratios of each treatment group were then divided by the mean of the ratios observed in matched control cells transduced with the NT1 or the GFP-expression vector to calculate the fold changes given in Table 2.

To measure the effect of candidate genes and Rev on the nuclear export efficacy (35) for HIV-1 US RNA for each treatment group, the mean US copy numbers/ng RNA extracted from the replicates of the cytoplasmic fractions were divided by the mean US copy numbers/ng RNA extracted from replicates of the whole cell lysates. The ratio of each treatment group were then again divided by the ratios observed in controls cells transduced with the NT1 or the GFP-expression vector to calculate the fold changes given in Table 2.

Fold changes for cytoplasmic levels of US or FS RNA were calculated by dividing the mean US or FS copy numbers per ng of extracted RNA of each treatment group by the mean derived from the control groups.

### RNA-seq

Cytoplasmic RNAs were extracted from the sorted BFP+ or unsorted control J-dual#3 cells with the Direct-zol RNA miniprep plus kit. RNA qualities were controlled by the 2100 Bioanalyzer instrument (Agilent) (data not shown). For each sample, a DNA fragment library was generated from the total RNA input using the Illumina TrueSeq Stranded mRNA kit and performed essentially as described by the manufacturer’s protocol (https://support.illumina.com/downloads/truseq-stranded-mrna-reference-guide-1000000040498.html). The libraries were sequenced on an Illumina HiSeq 2500 platform and the reads were saved together with a quality score. Raw data were converted to reads including a quality score by Software bcl2fastq v2.17. Sequences matching Illumina TrueSeq adapters were masked. Low quality bases, poly-A or poly-T stretches as well as masked bases were trimmed by Software fqtrim v0.9.5. Read quality was checked after sequencing and after base trimming by Software fastqc v0.11.8. For the viral genome, reads were aligned to HIV-Dual-GT using hisat2 (79). Read counting for the different features was performed using samtools (80) and regtools (81).

For splicing and differential expression analysis, reads were aligned to the human transcriptome using RSEM v1.2.20 (82) and Bowtie v1.1.2 (82) as read alignment program using default parameters. Differential expression analysis was performed by DESeq2 v1.18.1 (82) using the RSEM output for read counts per gene. We considered genes that showed an absolute log2-transformed fold change greater than 1 and a Padj value of less than 0.1 as differentially expressed. To exclude lowly expressed genes, we only considered those with mean TPM greater than 1 in either untreated or knockdown samples. Gencode v19 annotation was used in all steps of the analysis, including in the definition of protein coding genes. The reads mapping profiles were exhibited via the Integrative Genomics Viewer (IGV) v2.3 (83). Differential splicing analysis was performed with SUPPA (61) using TPM values from the RSEM transcript isoform output. For PSI calculation, events were required to pass the expression filter of TPM greater than 1 (-f 1 parameter). We considered splicing events with Padj values lower than 0.01 and absolute ΔPSI greater than 0.25 as differentially included or excluded. Polypyrimidine track length, and branch point scores were calculated using BPfinder (84). Splice-site strength was calculated using MaxENT (85) (hollywood.mit.edu/burgelab/software.html).

## Data availability

The raw and processed data of RNA-seq are accessible via the GEO (Gene Expression Omnibus) Accession Number GSE144347.

## Acknowledgements

H.X. was supported by a fellowship from the China Scholarship Council. The authors would like to acknowledge support from the Medical Facultýs core facilities “Cell Sorting and Immunomonitoring” and “Microarray / Next Generation Sequencing”. Plasmids psPAX2 and pMD2.G are gifts from Dr. Didier Trono. CRISPR reagents and libraries were a gift from Feng Zhang obtained from Addgene. Plasmid NL4-3-Δ6-drEGFP was kindly provided by Dr. Robert Siliciano.

## Author Contributions

K. Ü., A. E., B. G. and M.L. designed and supervised the study. H. X. and D. J. performed the experiments. H. X., E. W., M. M, P. K., A. B. O. V. S., A. E. and K. Ü. analyzed the data. H. X. and K. Ü. wrote the manuscript. A.E., B.G., E. W., M. M, P. K., A. E., A.B.O.V.S., M.L. and D.J. reviewed the manuscript.

## Supplementary figure legends

**Figure S1.**
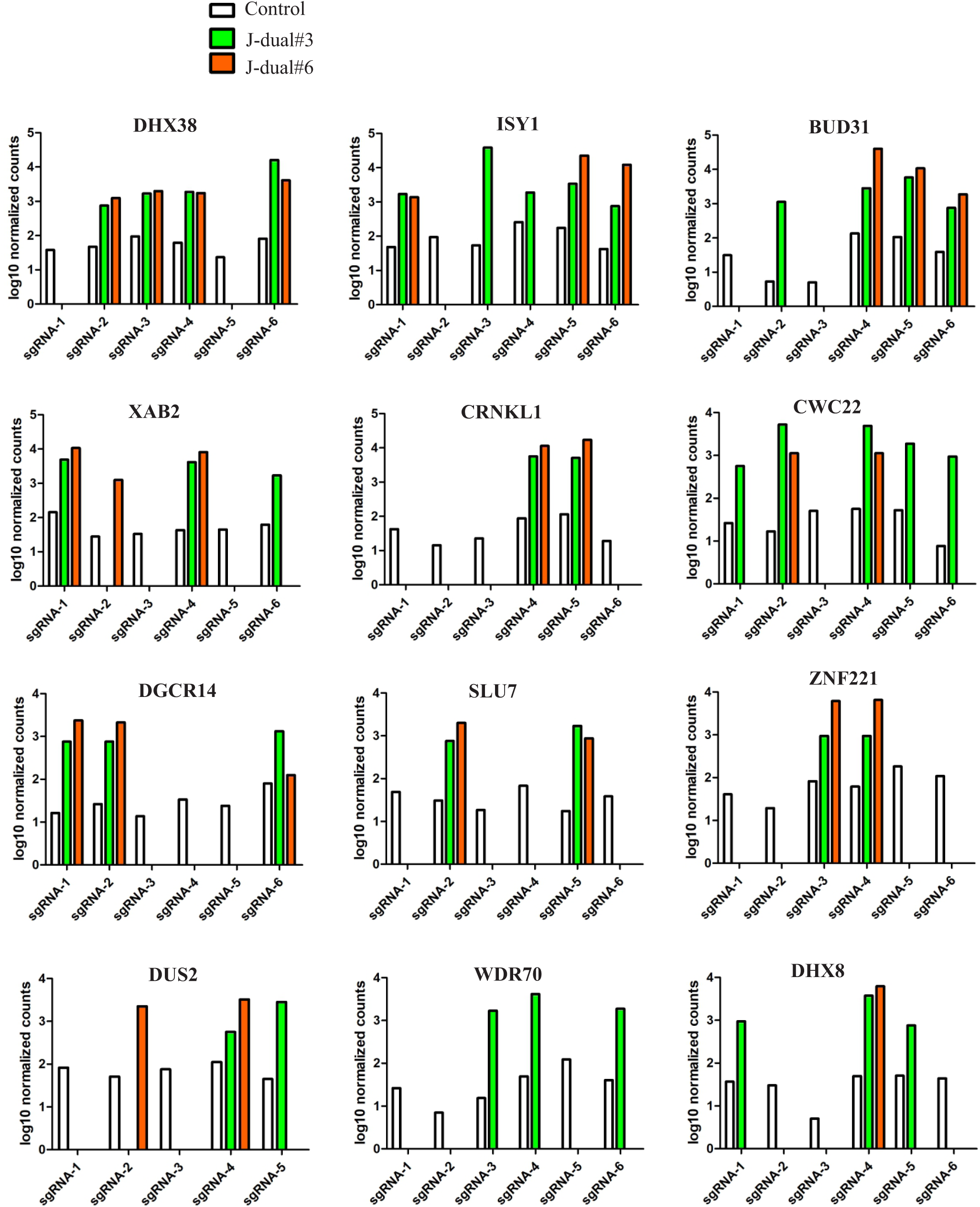
Enrichment of individual sgRNAs targeting the top 12 candidate genes. NGS data from unselected Jurkat cells (control) and selected J-dual#3 and J-dual#6 cells were analyzed by MAGeCK. Results are shown as normalized counts for each of the six guide RNAs targeting the indicated candidate genes.

**Figure S2.**
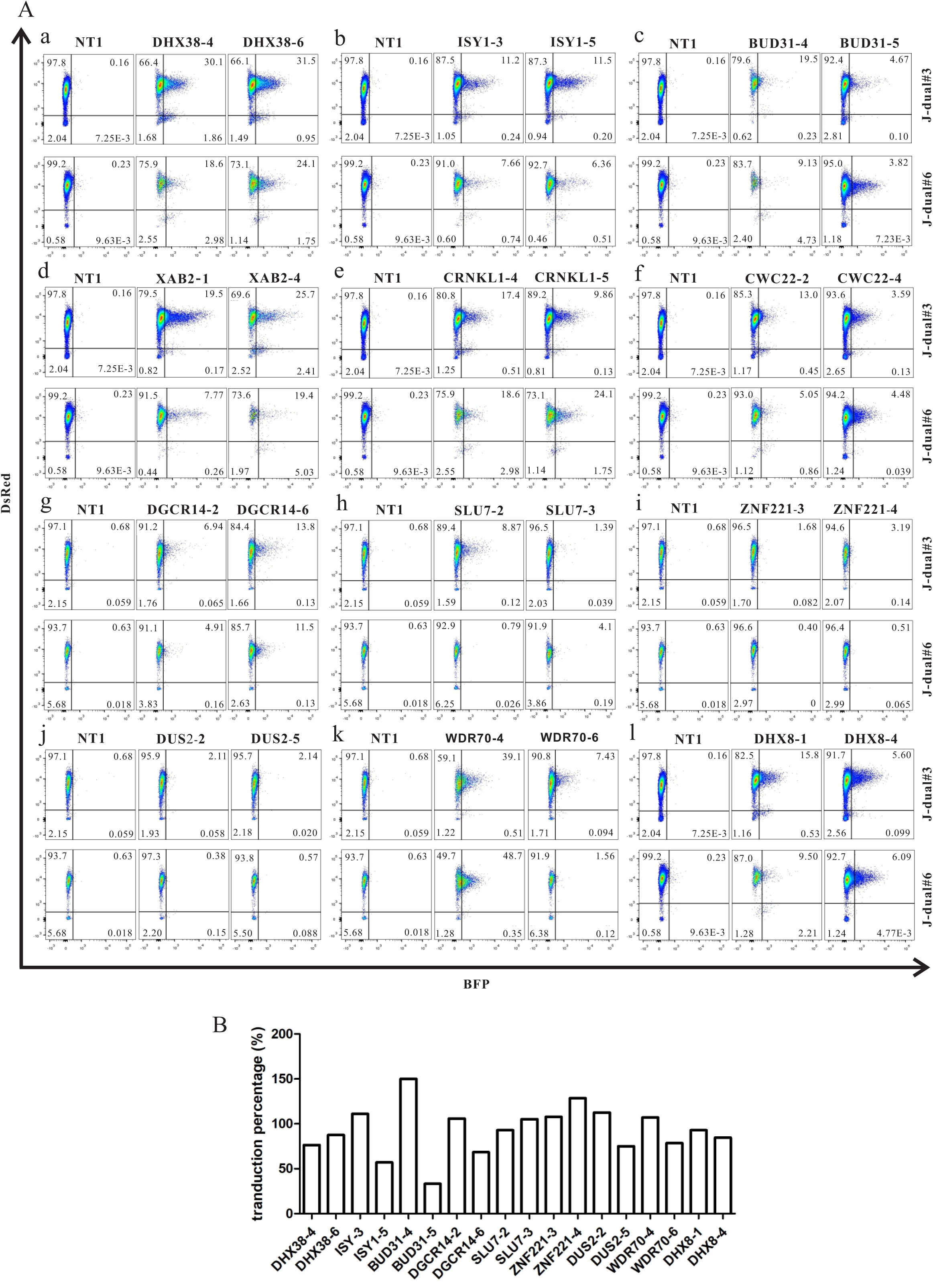
Validation of screening hits. (A) J-dual#3 and J-dual#6 reporter cells were co-infected with a lentiviral vector encoding Cas9 and a lentiviral vector expressing the indicated guide RNAs at an MOI of 1. Flow cytometric analyses were performed 7 days after infection. Each of the 12 candidate genes was targeted by two different sgRNAs. A non-targeting sgRNA (NT1) was used as a control. Representative results from at least 2 experiments are shown. (B) Percent transduction by individual lentiviral knock-out vectors. All individual lentiCRISPR-KO constructs were packaged into VSV-G pseudotyped lentiviral vector particles under the same conditions. The same volume of individual lentiCRISPR vector preparation was used to infect J-dual#3 cells. The percent transduction was calculated as described in Materials and Methods.

**Figure S3.**
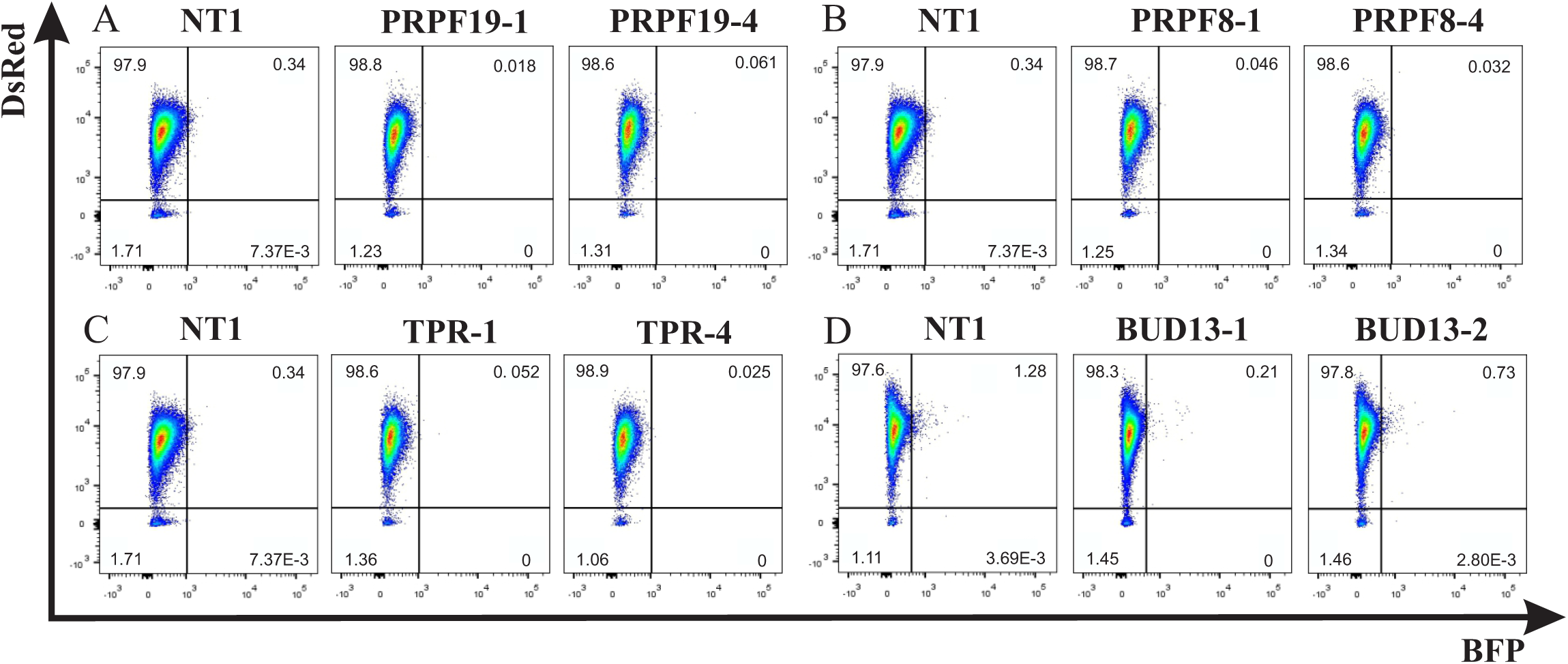
Influence of selected proteins reported to be involved in pre-mRNA splicing and nuclear retention on Gag-BFP expression. Two sgRNAs for each gene were picked from the GeCKO v2 library to target PRPF8, PRPF19, TPR and BUD13, respectively. The individual lentiCRISPR knockout vectors were constructed and used to transduce the J-dual#3 reporter cell line. Transduction efficiencies were confirmed (data not shown) and cells were analyzed by flow cytometry 7 days after transduction (A-D).

**Figure S4.**
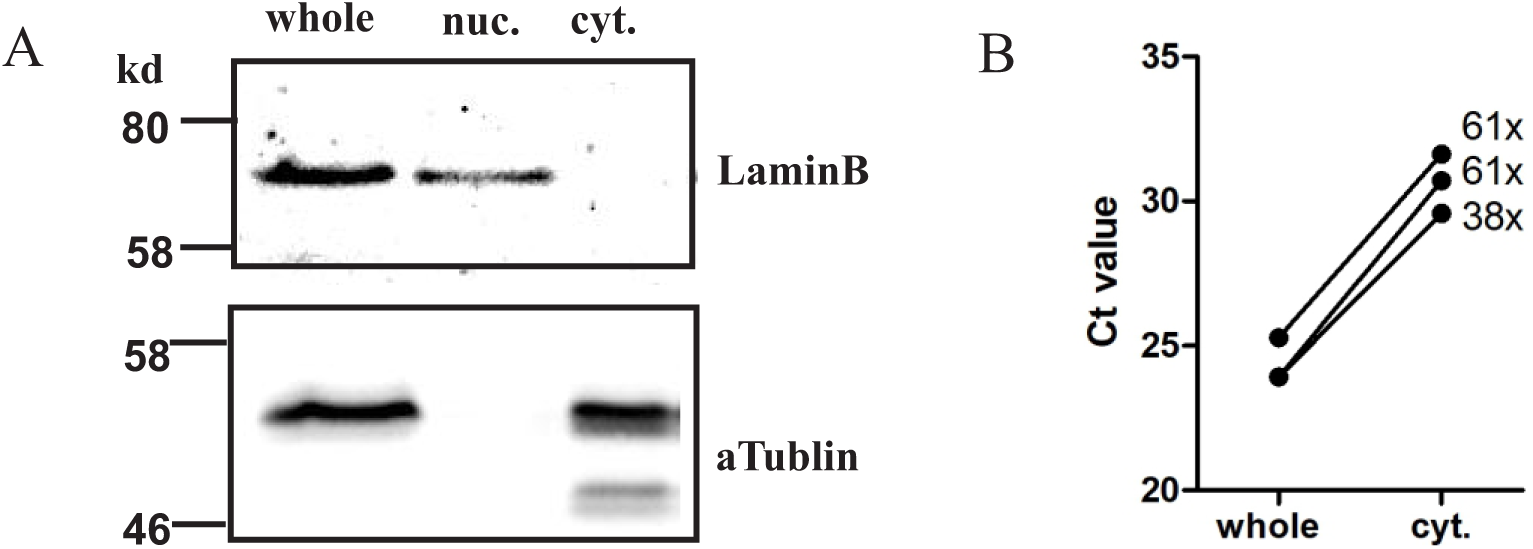
Validation of fractionation. (A) Representative Western blot analyses for LaminB (nuclear) and αTublin (cytoplasmic) in whole cell extracts and nuclear and cytoplasmic fractions from 1×10^6^ Jurkat cells. (B) Differences in GAPDH pre mRNA levels. CT values obtained by real time PCR from whole cell or cytoplasmic extracts from three experiments are shown. Numbers indicate the fold difference between GAPDH pre mRNA levels/weight of extracted RNA from whole cell and cytoplasmic extracts, assuming 2-fold differences in the targeted RNA copy numbers per ΔCT and adjusting for minor differences in the amounts of RNA added to the PCR reactions.

**Figure S5.**
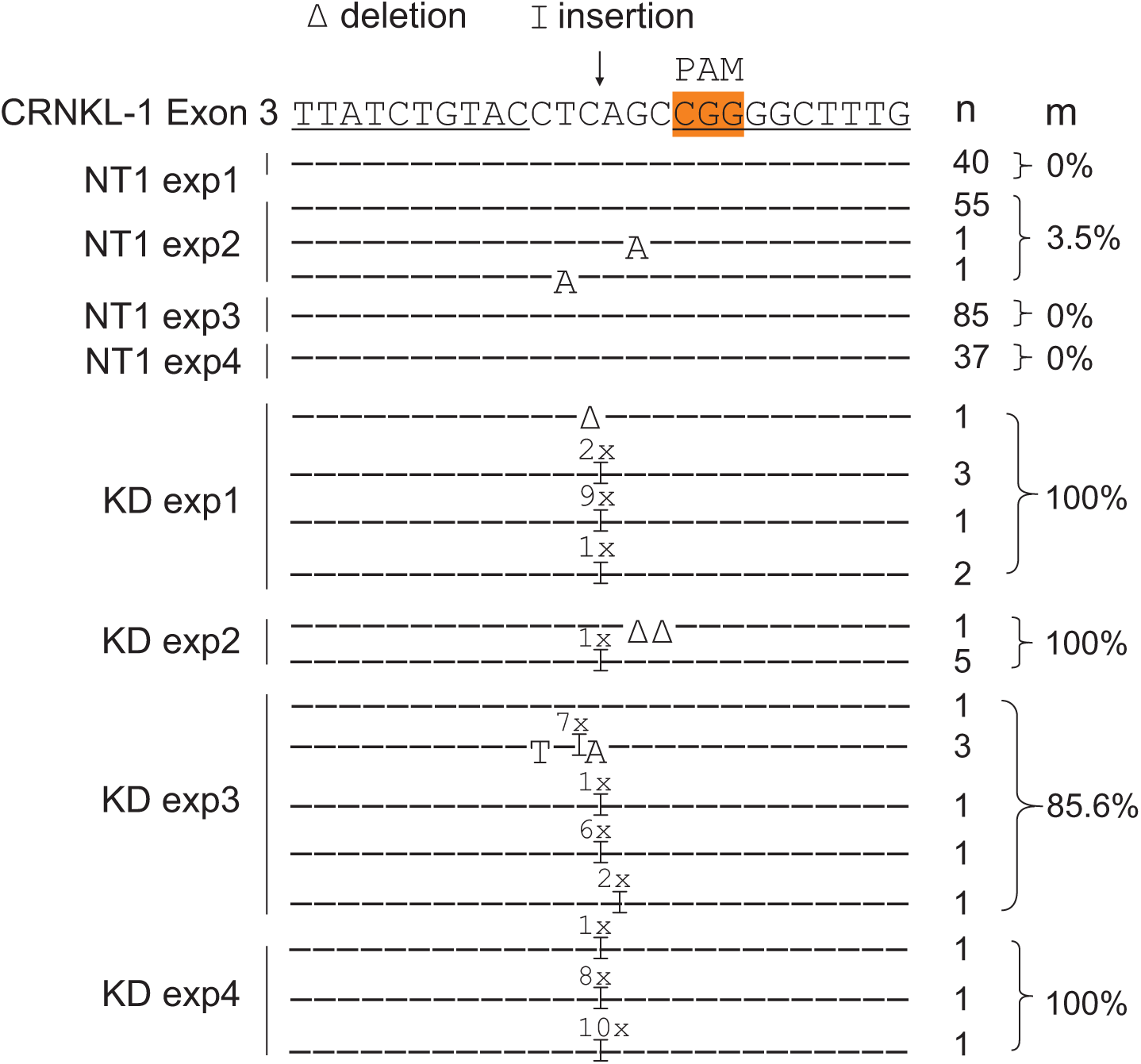
Number of reads and deviations from the wild type CRNKL1 target region after knock down of CRNKL1. The CRNKL1 sgRNA target region with the predicted Cas9 cleavage site (arrow) and the PAM region (shaded in orange) are aligned with reads from the transcriptome from four control (NT1 exp1-4) and four CRNKL1 knock down experiments (KD exp1-4). Underlined sequences were used to map reads to the target region. Point mutations, deletions (Δ) and insertions (Ι with number of nucleotides inserted given above) are indicated. “n”: number of reads with the respective sequence. “m”: percent mutation rate calculated as [100 x number of mutated reads / number of total reads mapped to the CRNKL1 target region].

**Figure S6.**
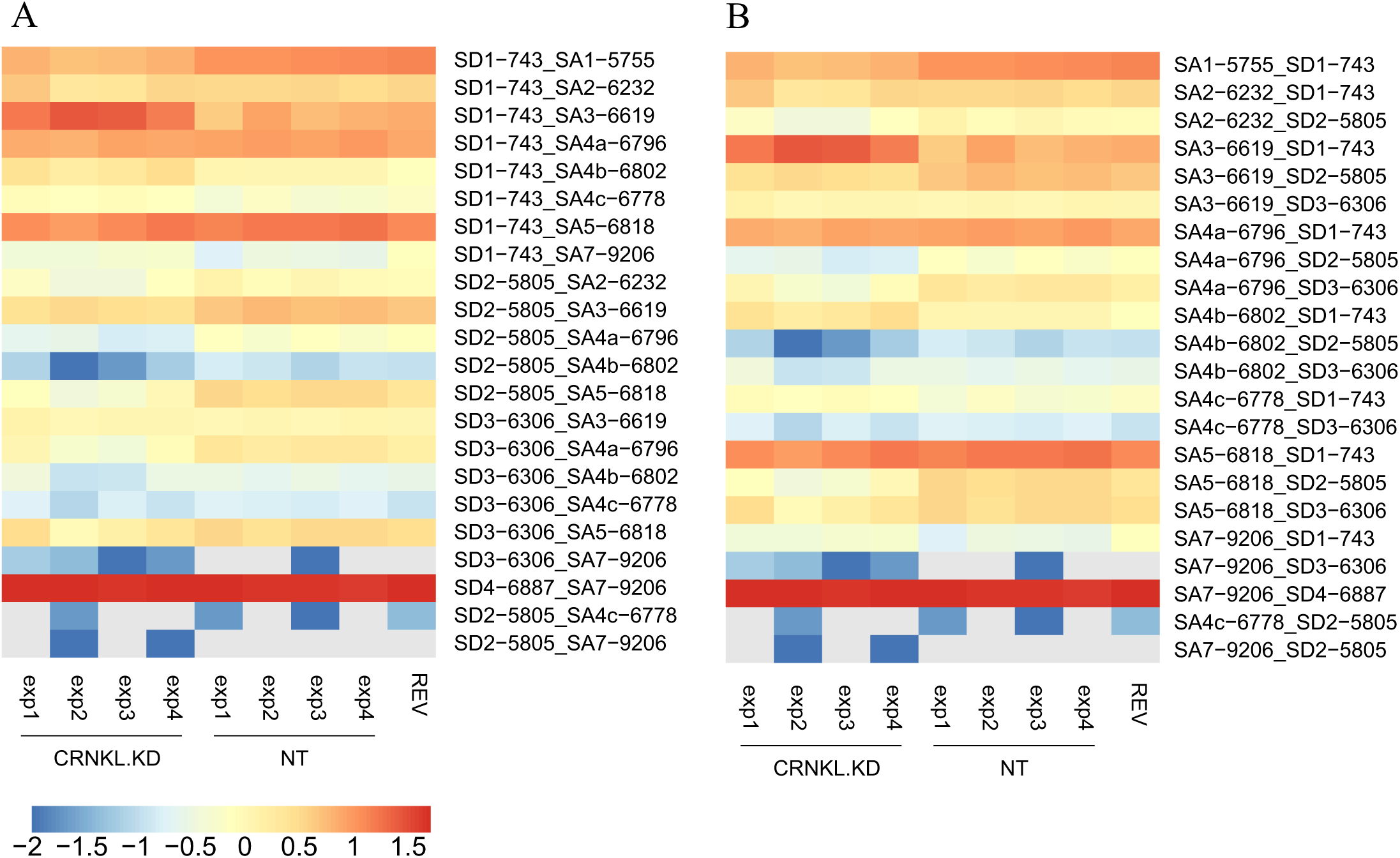
Changes in splice site usage of HIV RNAs upon CRNKL1 depletion. (A) Heat map representing, for every row, the relative frequency of indicated HIV-1 splice donor-splice acceptor pairs. Colors represent the log10 transformed percentages relative to all spliced HIV reads, as shown in the color bar underneath the heat map. Rows are sorted according to the splice donor sites. (B) As in A, but sorted for splice acceptor sites.

**Figure S7.**
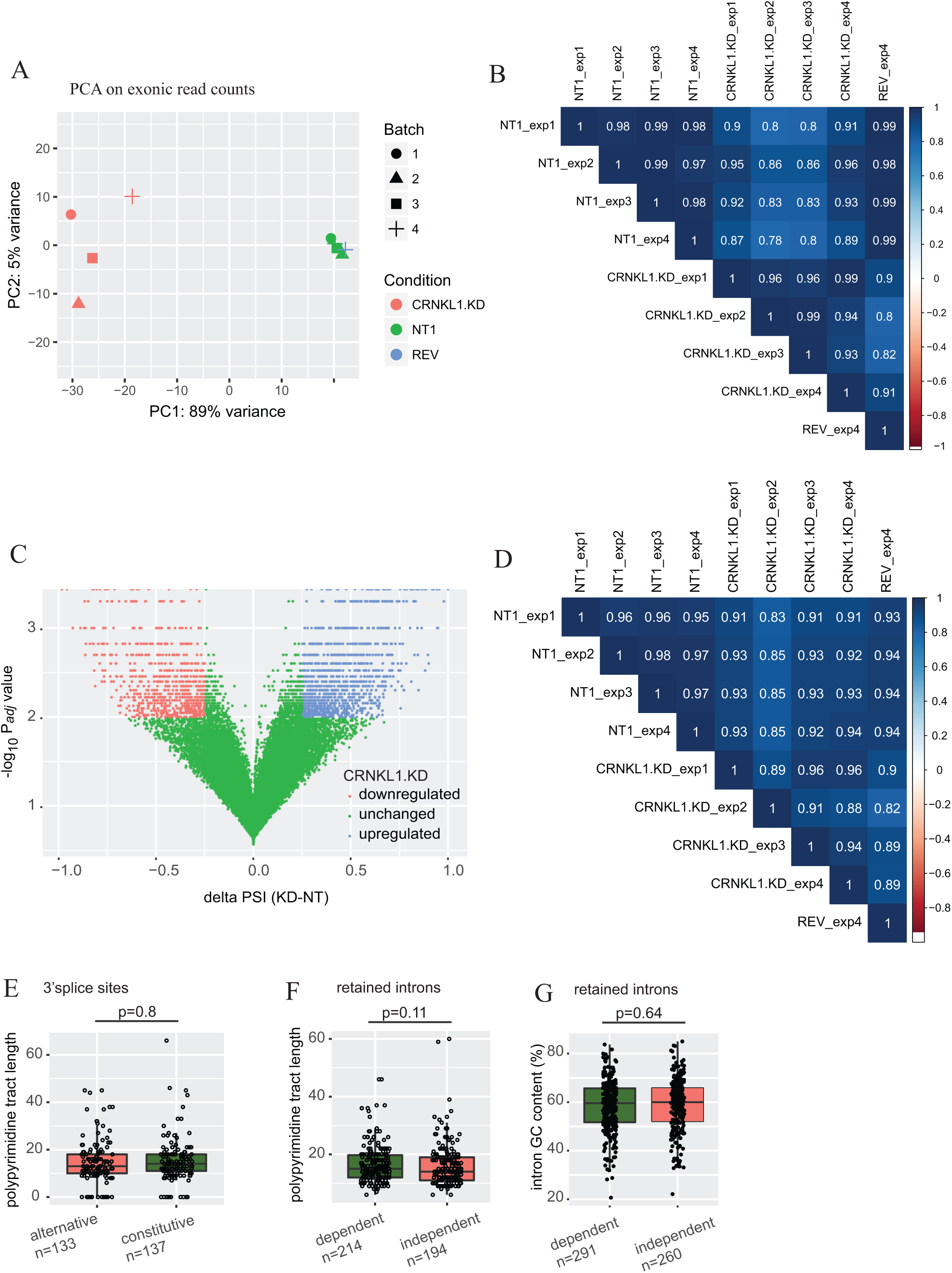
Quality control analyses for the transcriptomic data and additional analyses on differential splicing and intronic features. (A) Principal component analysis (PCA) of exonic read counts. Variance was assessed between biological replicates (1–4) and conditions (NT1, KD, REV). (B) Pairwise assessment of Pearson correlation coefficients between all exonic read counts from all RNA-seq samples. (C) Differential analysis of splicing events upon CRNKL1 KD. Volcano plot of ΔPSI (KD-NT1) values vs. negative log10-transformed Padj values is shown. To select up- and downregulated splicing events upon CRNKL1 KD (blue and red data points) the following cutoffs were used, respectively: ΔPSI (KD-NT1) >0.25, Padj <0.01; and ΔPSI (KD-NT1) <-0.25, Padj <0.01. (D) Pairwise assessment of Pearson correlation coefficients between all splicing event PSI values from all RNA-seq samples. (E) The length of polypyrimidine tracts of alternative 3’splice sites with higher usage under CRNKL1 KD condition were compared to those of constitutive 3’splice sites present in the same mRNAs. Wilcoxon rank sum test was used to test for significance. Absolute numbers of 3’splice sites in both groups are indicated. (F) Polypyrimidine track length was obtained by BPfinder for retained introns with higher inclusion upon KD (CRNKL1-dependent) and compared to that of unchanged retained introns upon KD (CRNKL1-independent), which were located in the same transcripts as CRNKL1-dependent ones. Box plots and data points are shown. Wilcoxon rank sum test was used to test for significance. Absolute numbers of retained introns in both groups are indicated. (G) GC content of retained introns with higher inclusion upon KD (CRNKL1-dependent) was compared to the GC content of unchanged introns upon KD (CRNKL1-independent), which were located in the same transcripts as CRNKL1-dependent ones. Box plots and data points are shown. Wilcoxon rank sum test was used to test for significance. Absolute numbers of retained introns in both groups are indicated.

**Figure S8.**
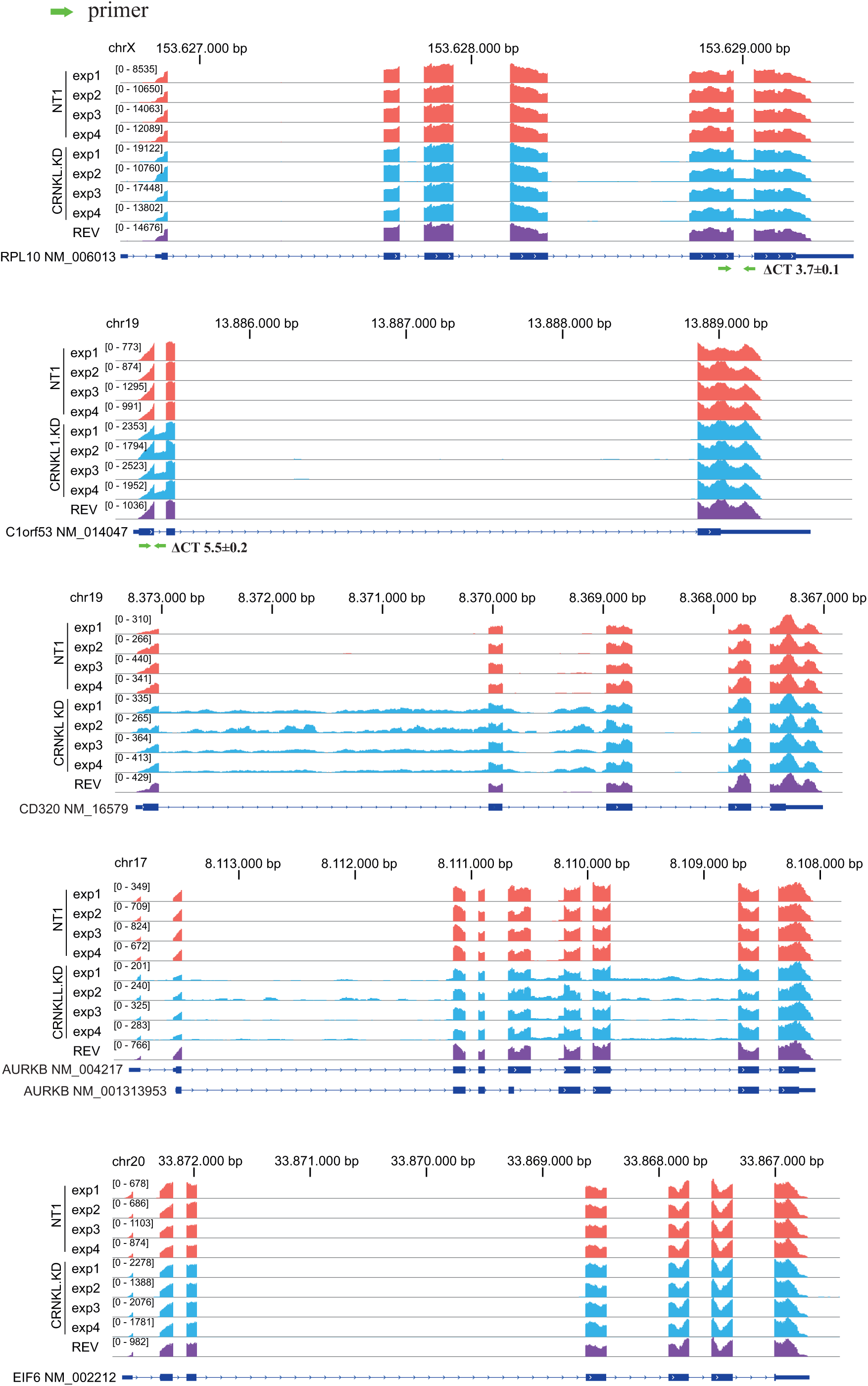
Examples and validation of intron retention in cellular transcripts after CRNKL1 knock-down. Mapping of the reads to genes with enhanced intron retention (RPL10, C19orf53 CD320), an alternative 5’and 3’splice site usage (5^th^ intron of AURKB) and an unaffected gene (EIF6) are shown for the four replicates of control (NT1) and CRNKL1 knock-down cells and the rev expressing cells. For RPL10 and C19orf53, intron retention was confirmed by RT-qPCR. Binding sites for specific primers spanning the exon-intron junctions are indicated by green arrows. RT-qPCRs were performed for relative quantification of respective intron-retaining transcripts using the cytoplasmic RNAs from the RNA-seq experiment. ΔCT(NT1-KD) ±SEM values obtained from the 4 replicates are indicated.

**Table S1.**
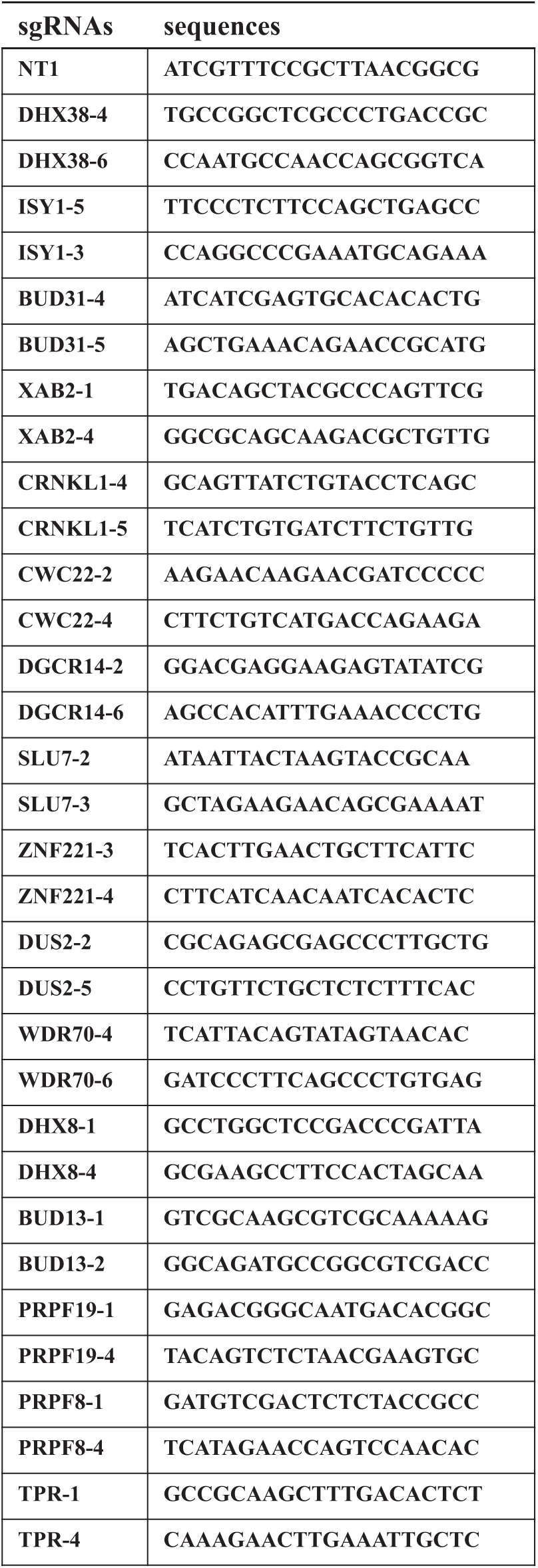
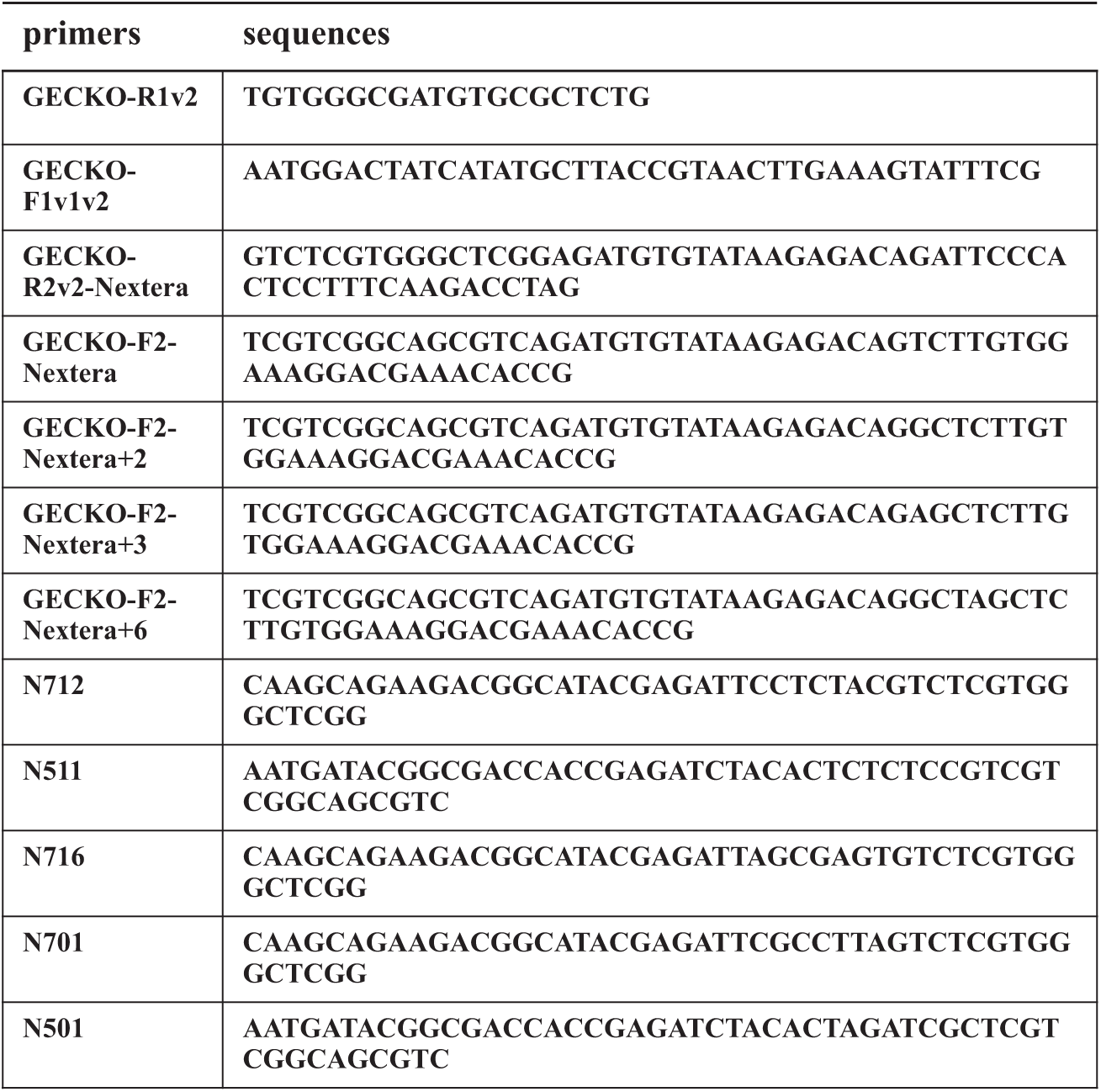
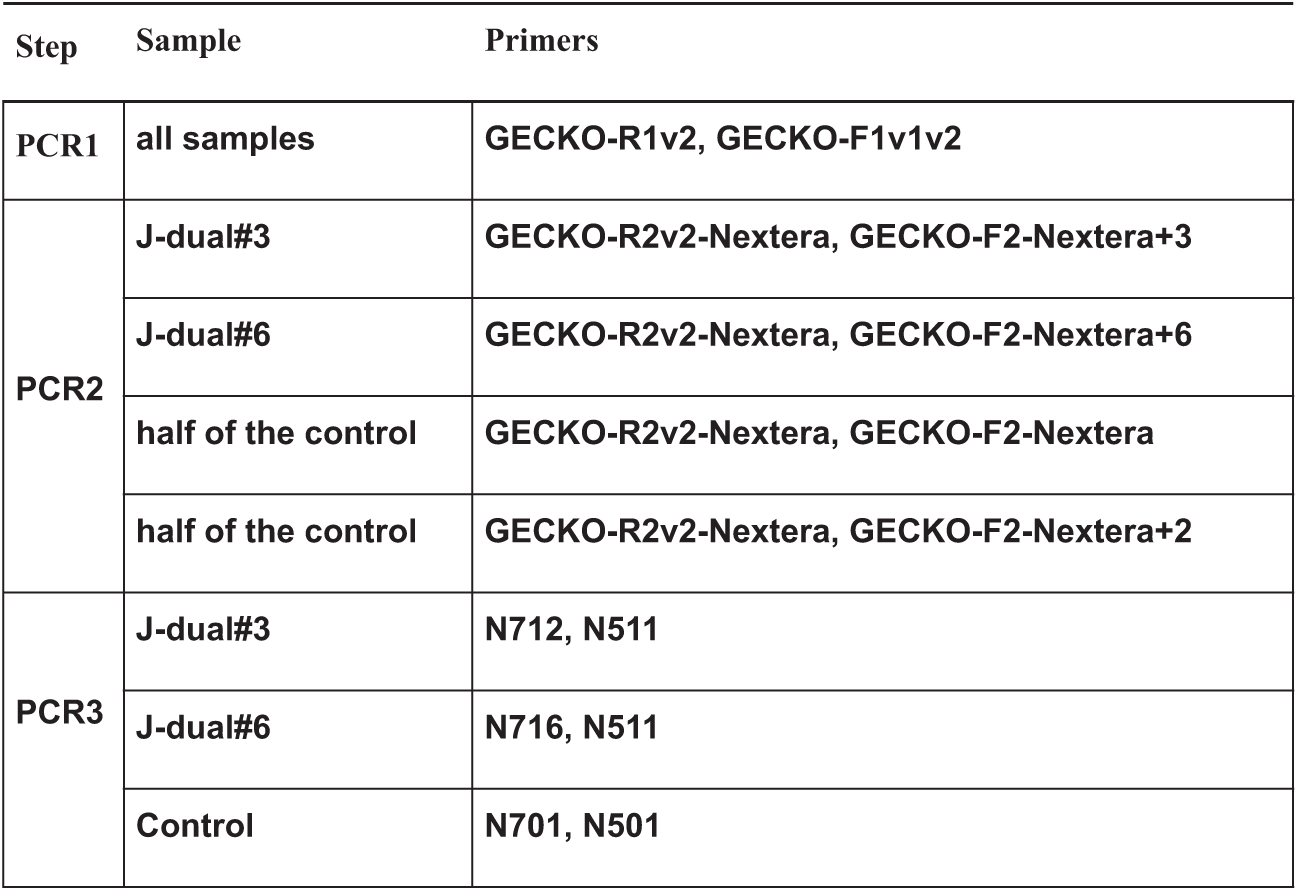
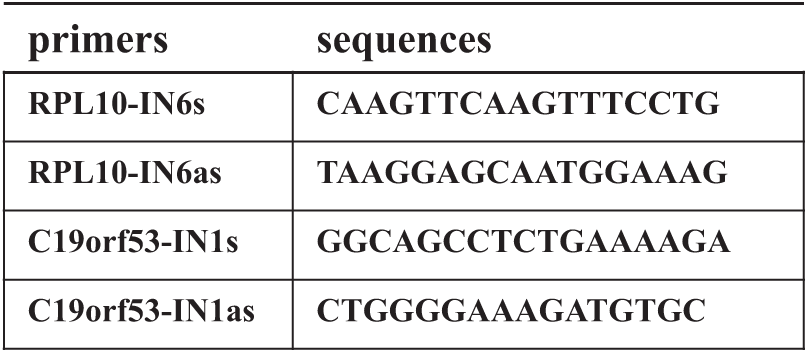
Primer sequences and combinations. (A) Sequences of sgRNAs used for screening hits validation. (B) Sequences of primers used for NGS libraries construction. (C) Primer combinations for NGS libraries construction. (D) Sequences of primers used for validation of retained introns.

